# On/off switches in the *DIVARICATA*-based regulatory network evolved through gene duplication, fusion, and truncation

**DOI:** 10.1101/2025.06.04.657860

**Authors:** Aniket Sengupta, Dianella Howarth

## Abstract

*DIVARICATA* (*DIV*) genes are *MYB* genes involved in many developmental processes in flowering plants. DIV-based regulatory networks involve paralogs of DIV (LFG and RADIALIS) that compete with DIV for either DNA or protein targets (like, the cofactor DRIF which is a distant paralog of DIV). These interactions act as on/off switches defining contrasting phenotypes. DIV proteins have multiple MYB domains; RADIALIS, LFG, and DRIF each have one. We determined the origin of these genes through Bayesian phylogenetic reconstruction. We report that *DIV* genes are a fusion of two simpler *MYB* genes. The first gene had two *MYB* domains (*MYBA–MYB1*), and the other gene had one *MYB* domain and a non-*MYB HDI* domain (*MYB2–HDI*). The ancestral *DIV* genes, hence, had four domains in the following sequence: *MYBA–MYB1–MYB2–HDI*. The *MYBA* domain was later lost, likely through a non-gradual truncation event leading to the following configuration in later-diverging *DIV*: *MYB1–MYB2– HDI*. The *MYBA* and *MYB2* domains were derived from the SHAQKY clade. The *MYB1* of *DIV* and the *MYBD* domains of *DRIF* were derived from the clade associated with the *SANT2* domain of *ZUO1/ZRF* genes. *LFG* genes evolved from *DIV* by truncation of the *MYB1* domain and the gain of repressor domains. We discuss how truncation of multi-domain DIV into RADIALIS and LFG was recruited towards on/off switches. Components of the DIV-based regulatory network, or their homologs, are present in a diversity of eukaryotes suggesting that their interaction may be ancestral to a large group of eukaryotes.

## Introduction

Morphological novelties are often not associated with genes created *de novo*, but with modifications and co-option of existing genes. *MYB* genes—first described in avian myeloblastosis virus (Klempnauer *et al*., 1982)—are a large super-family of protein-coding genes characterised by their conserved MYB domain. MYB proteins are ancient; they are present across eukaryotes and have diverse functions—for example, symmetry of flowers (Galego & Almeida, 2002; Corley *et al*., 2005) and telomere length regulation in vertebrates (Smogorzewska *et al*., 2000). MYB domains are usually classified into three categories based on conserved residues—the myb-type HTH (helix-turn-helix) domain, which typically binds DNA; the SANT (switching-defective protein 3, adaptor 2, nuclear receptor corepressor, and transcription factor IIIB) domain, which typically binds proteins; and the MYB-like domain that can be involved in either of these functions (Sigrist *et al*., 2013; ‘PDOC00037 in PROSITE’). The distinction between these three categories is not sharp and many predictive tools often annotate a given MYB domain with multiple categories.

*MYB* genes originated in an ancestral eukaryote (Lipsick, 1996; Yang *et al*., 2003; Jiang *et al*., 2004; Lv *et al*., 2013), which was likely a simple, unicellular organism; hence a function in flower symmetry cannot be the ancestral function for *MYB* genes. Therefore, *MYB* genes have been co-opted towards newer functions during the diversification of eukaryotes. The role of *MYB* genes in flower symmetry is best understood in the flowering plant *Antirrhinum majus*. The gene *A. majus DIVARICATA* (*AmDIV*), and its paralog *A. majus RADIALIS* (*AmRAD*), are involved in defining the opposite sides of the flower (Galego & Almeida, 2002; Corley *et al*., 2005). Close paralogs of *AmDIV* are also involved in other functions in flowering plants (angiosperms), such as in fruit development (Rose *et al*., 1999; Machemer *et al*., 2011; Song *et al*., 2023). AmDIV has two MYB domains—MYB1 and MYB2 (Fig. 1). The MYB1 domain is predicted to be a protein-binding SANT domain and the MYB2 domain is predicted variously by different prediction tools—as SANT (Lu *et al*., 2020), and as HTH myb-type (Sigrist *et al*., 2013; ‘DIVARICATA in UniProt’).

**Figure 1.**
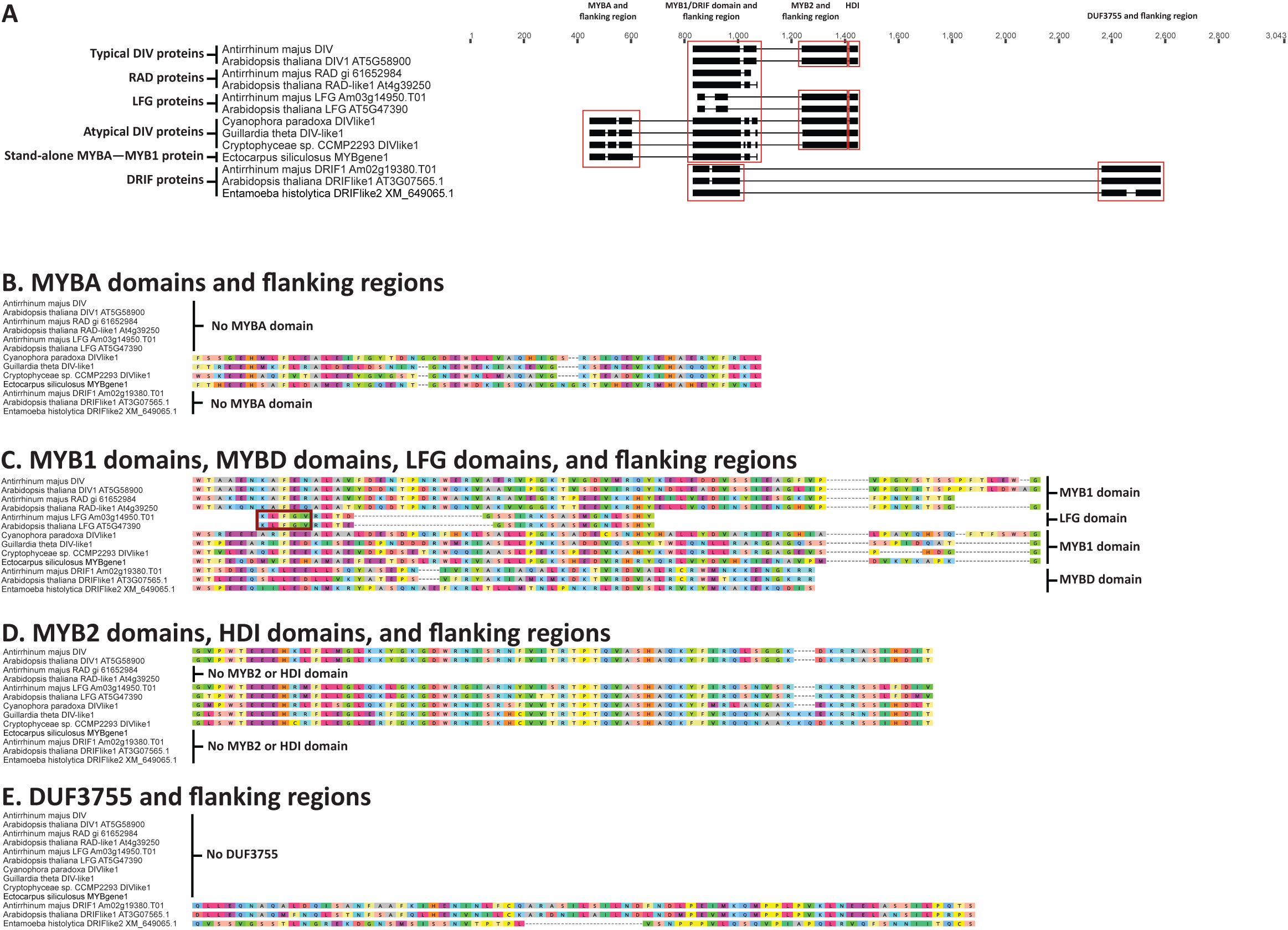
Alignment of the conserved domains of DIV, DRIF, RAD, and LFG homologs from representative eukaryotes. **A.** Amino acid alignment of DIV, DRIF, RAD, and LFG homologs. The conserved domains (MYBA, MYB1, MYB2, and HDI from DIV homologs, and the MYBD domain from DRIF) along with the conserved flanking sequences are displayed in dark boxes with lines representing gaps in the conserved region. Residues outside the conserved regions not marked by boxes or lines. **B.** Alignment of MYBA domains and flanking region. **C.** Alignment of MYB1 domains and the MYBD domain from DRIF along with the flanking region. Some Embryophyta proteins have limited conservation in this region, we describe them as LFG proteins based on the conserved Leucine-Phenylalanine-Glycine residues (in red box). LFG proteins also have other uniquely conserved residues (not shown). **D.** Alignment of MYB2 domains and flanking region, and the downstream HDI domain.

AmDIV functions as a transcription factor (Raimundo *et al*., 2013). It is likely that this DNA-binding activity of AmDIV is mediated via the MYB2 domain; this argument is based on indirect studies from truncated homologs—both artificial (Rose *et al*., 1999) and natural (Raimundo *et al*., 2013)—lacking the MYB2 domain. However, to be able to function, AmDIV needs to heteromerize with proteins called DIVARICATA AND RADIALIS INTERACTING FACTOR 1 and 2 (AmDRIF1 and AmDRIF2) (Raimundo *et al*., 2013) by likely employing its MYB1 domain. DRIF proteins have an atypical MYB domain (henceforth MYBD domain) and a downstream conserved region of unknown function called the DUF3755 (Raimundo *et al*., 2013). The MYBD domain of DRIF can bind to DIV proteins, at least in *Populus trichocarpa* (Petzold *et al*., 2018). The function of DUF3755 is not clear but in *P. trichocarpa*, this domain can bind to other transcription factors from the family WUSCHEL-RELATED HOMEOBOX (WOX) and KNOTTED1-LIKE HOMEOBOX (KNOX) (Petzold *et al*., 2018).

AmRAD acts as a competitive inhibitor of AmDIV. AmRAD has a single MYB1 domain, and it displays significant sequence identity with the corresponding MYB1 domain from AmDIV (Raimundo *et al*., 2013). AmRAD uses its MYB1 domain to bind with AmDRIF1 and AmDRIF2, thus inhibiting AmDIV from accessing these co-transcription factors (Raimundo *et al*., 2013). Thus, in *A. majus*, the competition between *AmDIV* and *AmRAD* act as an on/off switch defining two contrasting phenotypes. The activity of DIV is switched off by RAD by sequestering some of the co-factors DRIF1/2 to the cytoplasm, i.e., away from the nucleus where DIV is supposed to be upregulating genes (Raimundo *et al*., 2013). A similar, potentially orthologous (Sengupta & Hileman, 2022), DIV-RAD competition for DRIF has been reported in the fruit wall of tomatoes (*Solanum lycopersicum*) (Machemer *et al*., 2011).

A third on/off switch involving a DIV paralog is reported from rice (*Oryza sativa*) (Chen *et al*., 2019) where a DIV paralog called MYBS1 upregulates the gene *α-Amylase* (involved in starch hydrolysis) by binding to *cis*-regulatory sites (Chen *et al*., 2019). However, a MYB protein called MYBS2 that has only one MYB domain, can also bind to the same regulatory sites but represses *α-Amylase* (Chen *et al*., 2019). The serine residue at position 53 of MYBS2 is a phosphorylation site, whose phosphorylation is important for interaction with 14-3-3 proteins (Chen *et al*., 2019). The interaction between MYBS2 and 14-3-3 is likely involved in nucleocyctoplasmic shuttling of MYBS2 (Chen *et al*., 2019). The on/off switch in this network is also based on nucleocytoplasmic shuttling of proteins (Chen *et al*., 2019), similar to that of *A. majus* (Raimundo *et al*., 2013).

During sugar-starvation, the DIV paralog MYBS1 (that is localized in the nucleus) upregulates the transcription of *α-Amylase* (Chen *et al*., 2019). Under such conditions, MYBS2 is phosphorylated at serine-53, is bound to 14-3-3, and hence is sequestered in the cytoplasm, i.e., away from the gene *α-Amylase* (Chen *et al*., 2019). The activity of the DIV paralog MYBS1 is switched off by MYBS2 during sugar-abundance—the serine-53 gets de-phosphorylated, the 14-3-3 and MYBS2 do not interact, MYBS2 enters the nucleus and blocks the *cis*-regulatory sites of *α-Amylase* (Chen *et al*., 2019). Together, the DIV paralog MYBS1 and its competitor MYBS2 act as an on/off switch for starch hydrolysis.The origin of the multi-domain DIV proteins, or their paralogs involved in on/off switches has remained unexplained. The history of duplication, gain, or loss associated with the coming together of MYB1 and MYB2 domains in DIV proteins is not understood. There are many alternatives that may explain the presence of two MYB domains in the DIV proteins. First, an intragenic duplication (in which case one of the MYB domains from DIV should be nested within the other in a phylogenetic tree). Second, an intergenic translocation event (in which case the MYB1 and MYB2 clades should be reciprocally monophyletic in a phylogenetic tree). There are other potential events, such as, at least one of the domains arriving via horizontal gene transfer, and/or gene fusion. These hypotheses have remained untested.

The phylogenetic history of duplication and loss of *DIV* and *DRIF* genes has been examined in Viridiplantae (Green Plants) (Raimundo *et al*., 2018), however, their distribution outside this clade is not well studied. Further, how these individual MYB domains came to be a part of multi-domain genes is not understood and the origin of these domains is also not resolved. A close relationship between MYB1 and MYBD domains has been suggested but such nodes had a low support of 30–39% (Raimundo *et al*., 2018). To evaluate alternative scenarios explaining the origin of DIV homologs, we undertook a thorough survey of MYBA, MYB1, MYBD, and MYB2 (and HDI) domains in eukaryotes and then tested these hypotheses by employing phylogenetic methods. We surveyed all major clades of eukaryotes: Viridiplantae and their close relatives (Rhodophyta, Glaucophyta, and Cryptophyta), the next diverging clade of eukaryotes (Ochrophyta, Gyrista, Straminopila, and Haptophyta), and other distantly related eukaryotes (Unikonta and Excavata) (Supplementary Fig. 1). We determined the origin of the MYB domains involved in the DIV-based regulatory network and evaluated the molecular processes that may explain the loss of some of these domains. We interpret phylogenetic relationship among the MYB proteins involved in the DIV-based regulatory network to explain the origin of the on/off switches. We discuss the implications of these homologs co-occurring in the common ancestor of all or a large group of eukaryotes.

## Results and discussion

### Conserved domains in DIV and close homologs

The AmDIV protein has two MYB domains: MYB1 and MYB2 (Fig. 1). The MYB2 domain has the characteristic SHAQKY residues. There is another conserved region downstream of the MYB2 domain in AmDIV (between 3–7 residues downstream, depending on the boundary prediction for MYB2) with the sequence KDKRRASIHDIT (Fig. 1D). This is a highly conserved region across Viridiplantae and their close relatives with a majority-based consensus only varying from AmDIV in one amino acid. Previous workers have reported this conserved domain in flowering plants (described as motif 4 by Madrigal *et al*., 2019). We here describe this conserved region as the HDI domain, based on the HDI amino acids in this region. Hence, AmDIV has the following configuration: MYB1–MYB2–HDI. We describe such proteins with the configuration MYB1–MYB2–HDI as typical DIV. We find that the HDI domain is not limited to typical DIV proteins from flowering plants but is present downstream of MYB2 domains from a variety of eukaryotes (Fig. 2). All DIV proteins have MYB2 and HDI domains suggesting that the ancestral DIV protein also had MYB2 and HDI domains. There are other MYB proteins that also have MYB2 and HDI domains but not the MYB1 domain (that is always present in DIV proteins) (Fig. 2). Whether such proteins with MYB2 and HDI domains but no MYB1 domain are ancestral to DIV or derived from it through loss is discussed later.

**Figure 2.**
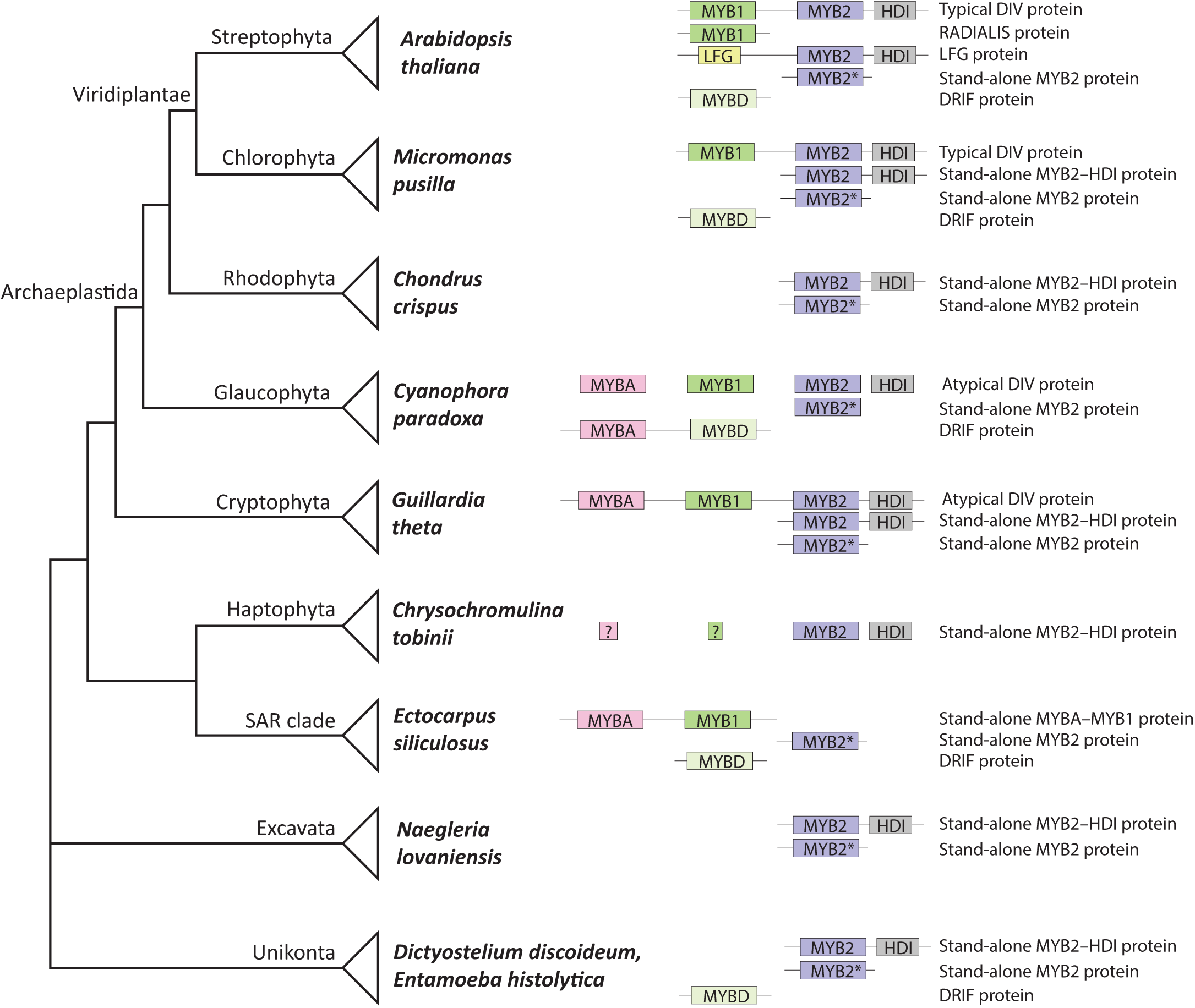
Distribution of DIV and DRIF homologs in representative eukaryotes. Relationships among the clades to which these species belong are represented by a simplified phylogeny of eukaryotes (adapted from Burki *et al*., 2012). Domain types are coded by names and color. MYB2* represents domains that are similar to the MYB2 domains of DIV proteins but do not have the downstream HDI domain. *Chrysochromulina tobinii* has a protein with clear MYB2 and HDI domains, and regions that have limited similarity with MYBA and MYB1 domains (NCBI conserved domain prediction tool predicts no MYB domain in these two regions). It is not clear whether *Dictyostelium discoideum* has a DRIF gene or not (potential homolog makes a polytomy with DRIF and other MYB), but *Entamoeba histolytica* does. The DRIF protein from *Cyanophora paradoxa* is unique in having a MYBA domain near the N-terminus in addition to the MYBD domain.

One such class of DIV homologs that have MYB2 and HDI domains but do not have a MYB1 domain can have some residues that weakly align with the MYB1 domains from other proteins. Examples of such proteins areAm03g14950.T01 in *Antirrhinum majus* (Fig. 1C) and the MYBS2 protein from rice that is involved in regulation of *α-Amylase* along with a DIV protein as a part of an on/off switch (Chen *et al*., 2019). These proteins have high sequence conservation upstream of the MYB2 domain, and also other conserved residues in the rest of the protein. We describe these proteins as LFG proteins based on the conserved LFG amino acid residues upstream of the MYB2 domain (Fig. 1C). The LFG domain has been called the R motif by other workers (Raimundo *et al*., 2018) but we describe them as the LFG domain to avoid a similarly named R domain from CYCLOIDEA proteins (Cubas *et al*., 1999) (DIV, DRIF, and CYCLOIDEA are a part of the same genetic network in *A. majus*) (Corley *et al*., 2005; Raimundo *et al*., 2013). The conserved LFG domain has sequence similarity to the plant-specific repression domain R/KLFGV (Ikeda & Ohme-Takagi, 2009). Thus, LFG proteins have the following domain configuration: LFG–MYB2–HDI. Other designations for LFG proteins have been suggested by previous workers—DIV-like or DVL (Raimundo *et al*., 2018)—but we will instead use the designation LFG proteins because “DIV-like” has been applied to a wide number of DIV homologs, whether they have a MYB1 domain or an LFG domain.

Some DIV homologs in algal groups diverging prior to Viridiplantae have an additional MYB domain upstream of the MYB1, for example, *Cyanophora paradoxa* DIV-like1 (Fig. 1A and B). Though reported in genome sequencing projects, this domain has not been studied in the context of evolution or function. We describe this domain as a MYBA domain. These algal DIV genes therefore have the following configuration: MYBA–MYB1–MYB2–HDI (Fig. 1A). We describe DIV paralogs with this configuration as atypical DIV, given that they have an additional domain.

There are some proteins, such as *Ectocarpus siliculosus* MYBgene1, that have a MYBA–MYB1 configuration, and do not have LFG, MYB2, or HDI domains (Fig. 1). There are some other proteins with simple configurations: proteins with only MYB2–HDI domains, proteins with only a MYB2 domain, and proteins with only a MYB1 domain (e.g., AmRAD).

It is important to note that though MYBA, MYB1, and MYB2 domains are present in distantly related organisms across eukaryotes, each of these domain types had a single origin. This is supported by the evidence that in a tree of MYB domains, the MYBA domains are monophyletic, and so are the MYB1 domains and the MYB2 domains (Supplementary Fig. 2). This suggests a single origin of DIV.

### Distribution of domains associated with DIV

To understand the evolution of DIV proteins—typical or atypical—we searched genomes and sequence databases for domains present in them and their close homologs (Fig. 2). Typical DIV, i.e., DIV with the configuration MYB1–MYB2–HDI, are found exclusively in Viridiplantae (Fig. 2). Viridiplantae, along with Rhodophyta and Glaucophyta, constitute the Archaeplastida (Fig. 1 and Supplementary Fig. 1). The immediate sister lineage of Viridiplantae, the Rhodophyta or Red Algae, do not have typical nor atypical DIV in any of the currently published genomes.

The two lineages that diverged prior to Rhodophyta are Glaucophyta and Cryptophyta, and these two lineages have atypical DIV (MYBA–MYB1–MYB2–HDI) (Fig. 2). This suggests the parsimonious scenario that atypical DIV with the configuration MYBA–MYB1–MYB2–HDI is ancestral to Archaeplastida+Cryptophyta, and that the MYBA domain was likely lost to give rise to the typical DIV seen in Viridiplantae. The corollary of this scenario is that DIV was lost in Rhodophyta, which is not surprising given that ancestral Rhodophyta underwent extensive genome reduction (Qiu *et al*., 2015). LFG proteins are present exclusively in the Streptophyta sub-clade of Viridiplantae (Fig. 2).

Stand-alone genes with the configuration *MYB2–HDI* are present in many members of Viridiplantae, Rhodophyta, Glaucophyta, Cryptophyta, Excavata, and Unikonta (Fig. 2), suggesting that this gene was likely inherited from the ancestral eukaryote. However, these proteins are not present in some eukaryotic lineages, like the SAR clade (Straminopila+Alveolata+Rhizaria) (Fig. 2). We also found domains that have high sequence similarity to the MYB2 domains of DIV, but which are not accompanied by any HDI domain (Fig. 2). Such MYB proteins with only MYB2 domains (marked as MYB2* in Fig. 2) are present across a wide diversity of eukaryotes, including the SAR clade. These are other MYB domains from the SHAQKY clade. It is possible that stand-alone *MYB2–HDI* genes are derived from stand-alone *MYB2* genes by acquisition of the *HDI* domain.

MYBA domains are found in one clade outside the Archaeplastida+Cryptophyta. They are present in the Gyrista, specifically, the Oomycota and Ochrophyta within Gyrista. Oomycota and Ochrophyta have stand-alone genes with the configuration *MYBA*–*MYB1* (Fig. 2). Gyrista is a clade within the larger SAR clade (Supplementary Fig. 1). However, other members of the SAR clade, outside Gyrista, do not have any MYBA or MYB1 domains. The Haptophyta species *Chrysochromulina tobinii* has a gene with *MYB2* and *HDI* domains, but it is not clear whether it has vestigial *MYBA* and *MYB1* domains (Fig. 2). Other members of Haptophyta that we searched do not have *DIV* genes or any stand-alone *MYB2–HDI* genes. There are no genes with only *MYBA* or only *MYB1* domains (except, the highly nested *RADIALIS* genes from seed-plants) in any eukaryote. Therefore, it is not clear what the ancestral genes with *MYBA* or *MYB1* domains looked like: whether they were stand-alone genes with only *MYBA* or only *MYB1* domains, or whether they were multi-domain genes. This is unlike *MYB2* domains, which is not only represented in *DIV* genes, but also in other genes with only *MYB2* domains (without any *MYB1* or *MYBA* domains).

Based on sequence similarity, the stand-alone *MYBA–MYB1* genes and stand-alone *MYB2–HDI* genes are related to the typical and atypical *DIV* genes. Given this varied domain organization among DIV homologs, and their uneven phylogenetic distribution, the following three hypotheses arise. First, did the genes with fewer domains arise from genes with more domains by loss; second, did the genes with fewer domains fuse to give rise to the genes with more domains; and third, is it a more complex history of both gain and loss. If *DIV* evolved from stand-alone *MYBA–MYB1* or stand-alone *MYB2–HDI* genes, then *DIV* should be nested within these genes in a phylogenetic tree. If stand-alone *MYBA–MYB1* genes and stand-alone *MYB2–HDI* genes evolved from *DIV* by loss of domains, these simpler genes should be nested within *DIV* in a phylogenetic tree. Phylogenetic trees have been used to distinguish between such hypotheses for similar questions (Yanai *et al*., 2002).

### *DIV* genes are derived from stand-alone *MYB2–HDI*

To understand the origin of *DIV* in relation to the stand-alone genes with the configuration *MYB2–HDI*, we reconstructed the phylogenetic history of genes that have both *MYB2* and *HDI* domains. This includes typical *DIV*, atypical *DIV*, stand-alone *MYB2–HDI* genes, and *LFG* genes (Fig. 3). We rooted the tree at its mid-point, given the challenges of determining the correct root among ancient lineages.

**Figure 3.**
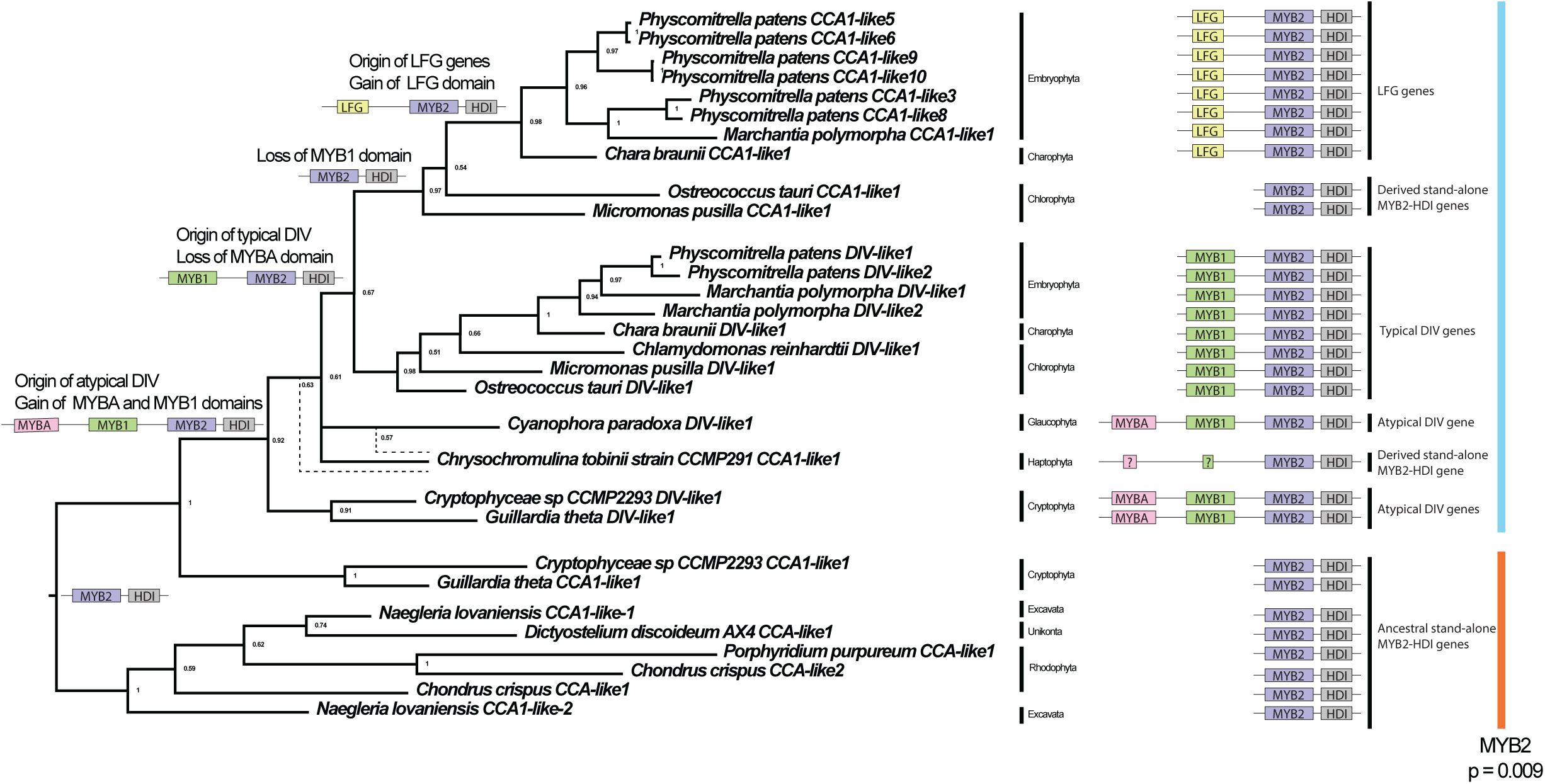
Nucleotide-based Bayesian phylogeny of genes that have *MYB2* and *HDI* domains. The tree was rooted at the midpoint. Bayesian posterior probabilities presented at nodes. Dotted lines represent alternative placements from different alignments. Domain organization represented on the right. Hypothesized ancestral states depicted at major nodes. Test of selection (relaxed vs. intensification) was performed on the *MYB2* domains of the groups designated with vertical blue bar (test group) and orange bar (reference group, i.e., *MYB2* domains from ancestral, stand-alone genes with the configuration *MYB2–HDI*). The *MYB2* domains from the test group are experiencing significant intensification of selection (p = 0.009, LR = 6.85, K = 1.08).

In our phylogenetic analysis *DIV* genes are nested within stand-alone *MYB2–HDI* genes providing evidence that *DIV* genes evolved from stand-alone *MYB2–HDI* genes by addition of *MYBA* and *MYB1* domains. *DIV* genes evolved at the base of Archaeplastida+Cryptophyta. The first *DIV* genes were atypical, that is, had a *MYBA* domain. Typical *DIV* genes evolved before the diversification of Viridiplantae by loss of the *MYBA* domain.

There are other loss and/or reductions associated with *DIV* genes. No genome from Rhodophyta that we surveyed had a copy of *DIV*. The gene *CCA1-like1* from the Haptophyta *Chrysochromulina tobinii* has a gene with *MYB2* and *HDI* domains but it is not clear whether it has no or vestigial *MYBA* and *MYB1* domains (Fig. 2), however is clearly nested within *DIV* (Fig. 3). In a tree of genes with *MYB2* and *HDI* domains, this gene defies the expectations from the species tree by being variously located (based on alternative alignments) close to Glaucophyta and Cryptophyta but with poor support (Fig. 3).

We searched the genomes of other Haptophyta species in GenBank (*Isochrysis galbana* GCA_018136815.1, *Emiliana huxleyi* GCA_000372725.1, *Diacronema lutheri* GCA_019448385.1, *Chrysochromulina parva* GCA_002887195.1, Pavlovales sp.

GCA_026770615.1), but none of them, including another species from the same genus *Chrysochromulina parva*, has a copy of this gene. This suggests that the gene *CCA1-like1* from *Chrysochromulina tobinii* was likely acquired by horizontal gene transfer from a Cryptophyta or Archaeplastida species followed by a loss of the *MYBA* and *MYB1* domains.

### *DIV* genes are derived from stand-alone *MYBA–MYB1*

To understand the origin of *DIV* in relation to the stand-alone genes with the configuration *MYBA–MYB1*, we reconstructed the phylogenetic history of genes that have *MYBA* and/or *MYB1* domains. This includes typical *DIV*, atypical *DIV*, and stand-alone *MYBA–MYB1* genes (Fig. 4). We rooted the tree at its mid-point.

**Figure 4.**
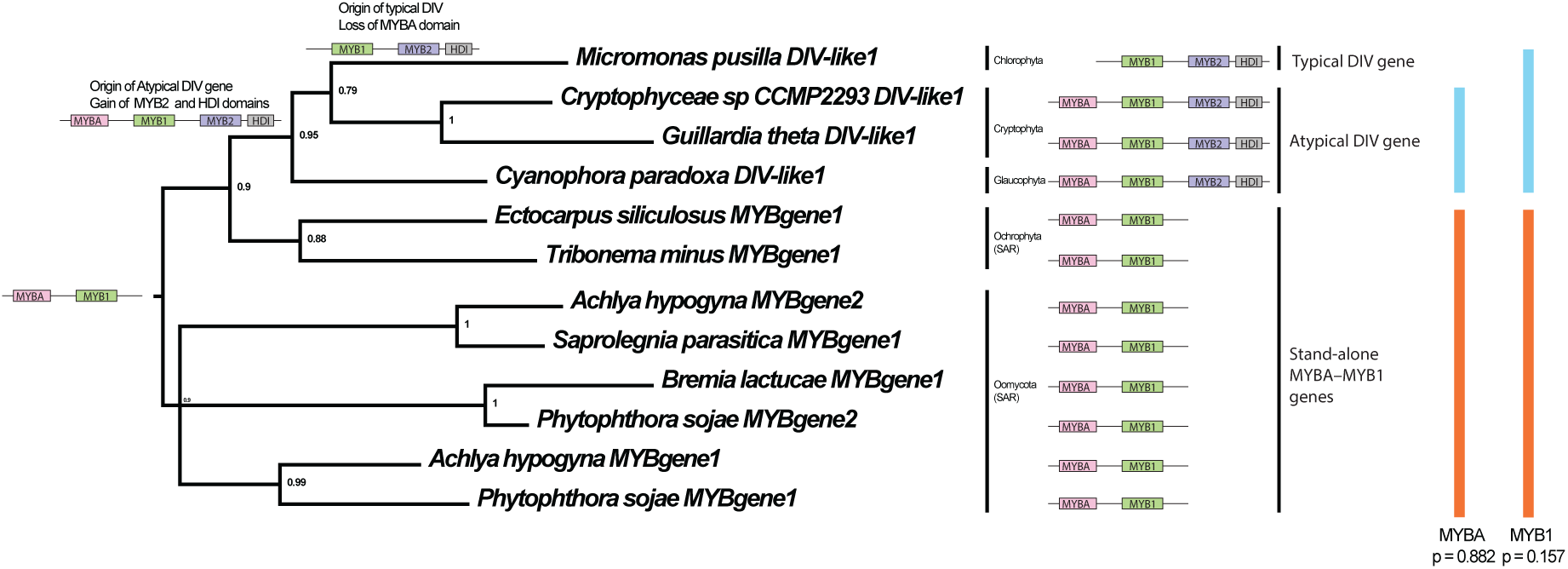
Nucleotide-based Bayesian phylogeny of genes that have *MYBA* and *MYB1* domains. The tree was rooted at the midpoint. Bayesian posterior probabilities presented at nodes. Domain organization represented on the right. Hypothesized ancestral states depicted at major nodes. Test of selection (relaxed vs. intensification) was performed on the *MYBA* domains of the groups designated with vertical blue bar (test group) and orange bar (reference group, i.e., *MYBA* domains from ancestral, stand-alone genes with the configuration *MYBA–MYB1*). A similar test was performed on *MYB1* domains. In both cases, there is no significant difference between the test group and the reference groups in terms of relaxed or intensified selection. *MYBA*: p = 0.882, LR = 0.02, K = 0.99; *MYB1*: p = 0.157, LR = 2.00, K = 0.63. *Micromonas pusilla DIV-like1* was not included in the test of selection for *MYBA* because it has no *MYBA* domain.

*DIV* are nested within stand-alone *MYBA–MYB1* genes providing evidence that *DIV* genes evolved from stand-alone *MYBA–MYB1* genes by addition of *MYB2* and *HDI* domains (Fig. 4). *DIV* genes evolved at the base of Archaeplastida+Cryptophyta (Fig. 4). The first *DIV* genes were atypical, that is, had a *MYBA* domain (Fig. 4). Typical *DIV* genes evolved before the diversification of Viridiplantae by loss of the *MYBA* domain (Fig. 4).

*DIV* are nested within stand-alone *MYBA–MYB1* and stand-alone *MYB2–HDI* genes, suggesting that *DIV* evolved from these stand-alone genes. But how did the two separate clades of genes come together?

### *DIV* genes evolved by gene fusion

The first *DIV* genes were atypical and had conserved domains in the following order: *MYBA– MYB1–MYB2–HDI*. *DIV* genes are nested within the stand-alone genes, and the order of the domains are the same as the stand-alone genes. This suggests that the first *DIV* genes originated by the fusion of a stand-alone *MYBA–MYB1* gene with a stand-alone *MYB2–HDI* gene. During the fusion the order was such that *MYBA–MYB1* was at the 5′ end and *MYB2–HDI* at the 3′ end.

One question that immediately arises is about the source of the contributing stand-alone genes. Did they come from existing genes in the same genome, or from another species through horizontal gene transfer? The following line of evidence suggests that the contributing stand-alone *MYB2–HDI* gene came from the same genome where the first *DIV* gene evolved, as opposed to by horizontal gene transfer. *DIV* evolved in the common ancestor of Archaeplastida+Cryptophyta, and both clades have ancestral, stand-alone *MYB2–HDI* genes (at least in Rhodophyta and Cryptophyta) (Fig. 3). Furthermore, the closest relatives of *DIV* in a tree of genes with *MYB2–HDI* domains are the stand-alone *MYB2–HDI* genes from Cryptophyta (Fig. 3). This suggests that the hypothetical stand-alone *MYB2–HDI* gene that was sister to the Cryptophyta stand-alone *MYB2–HDI* genes (represented in Fig. 3 by *Guillardia theta CCA1-like1* and *Cryptophyceae sp. CCMP2293 CCA1-like1*) became the *DIV* gene by fusing with a stand-alone *MYBA–MYB1* gene.

This elicits the questions about the source of this contributing *MYBA–MYB1* gene that are found only in Gyrista as stand-alone genes. Neither Cryptophyta nor Archaeplastida have stand-alone *MYBA–MYB1* genes. However, Cryptophyta and the early-diverging Archaeplastida lineage Glaucophyta have *MYBA–MYB1–MYB2–HDI* genes. Did the stand-alone *MYBA–MYB1* gene that participated in gene fusion come through horizontal gene transfer from Gyrista to the ancestor of Archaeplastida+Cryptophyta? It is important to note that there have been several independent eukaryote-to-eukaryote endosymbiosis events (Keeling, 2010). These endosymbiosis events, along with non-endosymbiosis associated horizontal gene transfer, make it difficult to determine the species where a gene first appeared. There have been proposals that the common ancestor of all Stramenopila (including Oomycota and Ochrophyta was endosymbiotic with a Rhodophyta, but this has been challenged in more recent studies (Wang *et al*., 2017). If indeed the stand-alone *MYBA–MYB1* genes in Oomycota and Ochrophyta are derived from Rhodophyta through such an endosymbiotic event, then one would expect that at least some Rhodophyta species would have a similar gene—but that is not the case because no Rhodophyta genome available in GenBank has a stand-alone *MYBA–MYB1* gene. Indeed, recent studies caution us to consider not only endosymbiosis but also the possibility of horizontal gene transfer or ancient origin (followed by repeated loss) when comparing genes between Oomycota and Archaeplastida (Wang *et al*., 2017).

We analysed the scenario of ancient origin (followed by repeated loss) in which the contributing *MYBA–MYB1* was present in the common ancestor of Gyrista and Cryptophyta+Archaeplastida. In this case, the native stand-alone *MYBA-MYB1* gene would have fused with the native stand-alone *MYB2–HDI* gene in the common ancestor of Cryptophyta+Archaeplastida. This scenario would involve several independent losses of stand-alone *MYBA-MYB1* genes across a diverse group of eukaryotes, but one may expect more than just two distant lineages (Gyrista and Cryptophyta+Archaeplastida) to retain the *MYBA* and *MYB1* domains. The other possibility is that the common ancestor of Cryptophyta+Archaeplastida acquired a stand-alone *MYBA–MYB1* gene from Gyrista through horizontal gene transfer (for an example of horizontal gene transfer followed by fusion, see Djuika *et al*. 2015). There is some, but limited, phylogenetic evidence for this scenario for *DIV* genes: in a tree of genes with *MYBA* and/or *MYB1* domains, *DIV* genes are nested within genes from Gyrista (Fig. 4). The closest relatives of *DIV* genes among Gyrista are stand-alone *MYBA-MYB1* from Ochrophyta (represented by species from Phaeophyta and Xanthophyta in our tree) (Fig. 4). However, evaluation of these two scenarios—ancient origin versus horizontal gene transfer—would require large scale phylogenomic and syntenic analyses across all major clades of eukaryotes.

Whatever the source of the contributing stand-alone genes, it is likely that a fusion event generated the first *DIV* gene. Fusion between a *MYBA–MYB1* gene and a *MYB2–HDI* gene would have given rise to an atypical *DIV* gene. Such *DIV* genes are present in Cryptophyta and Glaucophyta. However, the *DIV* genes in Viridiplantae, including *AmDIV*, lack the *MYBA* domain, suggesting that the *MYBA* domain was lost before the diversification of Viridiplantae.

### The MYBA domain was lost through a non-gradual process

If a conserved domain is absent in later diverging lineages, then two underlying processes may explain such a loss. First, the domain may have been lost through a saltatory event, like a segmental deletion. Secondly, it may have been lost through a gradual degenerative process.

MYBA and MYB2 domains are from the same clade of MYB domains (Supplementary Fig. 2), and presumably can have similar functions. If they co-occur in a protein, it is possible that one of the domains would be lost due to redundancy. This scenario would be comparable to the gradual pseudogenization of a paralog after duplication of a gene (example of duplicates on route to being pseudogenes in McGrath *et al*., 2014; Lehti-Shiu *et al*., 2015).

When a protein coding region is lost through a degenerative process, like pseudogenization, the following two lines of evidence can manifest. First, each subsequent branch in the gene tree may display increased pseudogenization/degeneration until all signs of homology are lost. Second, the redundant region may display relaxation of selective pressure relative to the paralog that was retained.

We tested these hypotheses in the context of the loss of the *MYBA* domain. We searched the *DIV* genes and their upstream regions from early-diverging Viridiplantae for signs of the *MYBA* domain or its vestiges. We did not find any evidence of vestigial *MYBA* domains in the tested regions. Further, because the *MYBA* domain is located near the start codon, even one non-sense or frameshift mutation could have pulled the remainder of the gene into pseudogenization—but *DIV* genes in Viridiplantae are not pseudogenized. Though the assessment may change as newer genomes become available, currently we find no support for the hypothesis that early-diverging Viridiplantae have vestigial *MYBA* domains in various stages of degeneration.

Next, we determined whether the *MYBA*, *MYB1*, and *MYB2* domains (including some conserved flanking regions) are undergoing a relaxation of selective pressure after the fusion event (Fig. 3 and Fig. 4). Selection pressure did not relax on any of the three *MYB* domains after fusion relative to the domains in the pre-fusion, ancestral stand-alone genes. In fact, the *MYB2* domain is undergoing an intensification of selection after the fusion event (Fig. 3).

The lack of relaxed selection on *MYBA* domains after gene fusion (Fig. 4) and the absence of gradually degenerating, vestigial *MYBA* domains in early-diverging Viridiplantae (Fig. 4) suggests that the loss of the *MYBA* domain (associated with typical *DIV* genes) did not happen through a gradual degenerative process. Therefore, the loss of the *MYBA* domain likely happened through a saltatory event, like, an in-frame segmental deletion. This is further supported by the fact that the region corresponding to the deletion (between the starting methionine and the MYB1 domain) is shorter in typical DIV and also has many gaps when aligned with atypical DIV.

### LFG proteins evolved by truncation of typical DIV

A lineage of *DIV* in Viridiplantae lost the *MYB1* domain to secondarily become a stand-alone *MYB2–HDI* gene. Such genes are represented in Chlorophyta by *Micromonas pusilla CCA1-like1* and *Ostreococcus tauri CCA1-like1* (Fig. 3). The upstream regions of these two genes have no vestigial *MYB1* domains, suggesting that the truncation was likely a saltatory event, like a segmental deletion. The line of such secondarily evolved stand-alone *MYB2–HDI* genes in Streptophyta (*Chara*+Embryophyta in Fig. 3) underwent further modification to become *LFG* genes (Fig. 3). A phylogeny and alignment of *LFG* genes is available online in the Panther database (‘PTHR44191 in Panther’). The earliest diverging species with *LFG* genes that can be identified by BLAST searches are from *Mesostigma viride* and *Chlorokybus atmophyticus*, which are also the first diverging species in the Streptophycean line leading to Embryophyta (Wang *et al*., 2020). Thus, *LFG* genes are truncated *DIV* genes that evolved at the base of Streptophyta through the loss of the *MYB1* domain and the gain of some unique sequence features. Some characteristic conservations that evolved in the LFG proteins are the CCHC-type zinc finger domain, the LFG domain, a conserved phosphorylation site (serine residue), and the EAR domain.

LFG proteins usually have a conserved, predicted CCHC-type zinc finger domain at the N-terminus (this study and Huang *et al*.(2015)). CCHC-type zinc finger domains have diverse functions, including binding to nucleic acids (reviewed in Aceituno-Valenzuela *et al*., 2020; reviewed in Sun *et al*., 2022). The LFG domain is located immediately downstream of the zinc finger domain. LFG domains have sequence similarity to the plant-specific repression domain R/KLFGV (Ikeda & Ohme-Takagi, 2009). The serine residue at position 53 of MYBS2 in rice (corresponding to position 41 in *A. majus* protein Am03g14950) is conserved across LFG proteins. In rice, this serine residue is a phosphorylation site, whose phosphorylation is important for interaction with 14-3-3 proteins (Chen *et al*., 2019); this interaction has been predicted to be involved in nucleocyctoplasmic shuttling of MYBS2 important for switching off the function of the DIV paralog MYBS1 (Chen *et al*., 2019)

Several LFG proteins have an additional predicted repressor domain near the C-terminus called the EAR motif; such motifs have the characteristic sequence LXLXL (Chow *et al*., 2023). The EAR motif of the LFG protein AT5G47390 (present in *A. thaliana*) has been characterized as a transcriptional repressor (Huang *et al*., 2015). EAR motifs are present in many, though not all, LFG proteins, including the one from an early diverging Streptophyte, *C. atmophyticus*. This suggests that the EAR domain is possibly ancestral to LFG proteins, but has been lost several times.

LFG proteins retain their DNA-binding MYB2 domains. We hypothesize that the first LFG proteins acted as competitive inhibitors of their closes relatives from which they were derived by the loss of the MYB1 domain. These LFG proteins would continue to bind to the *cis-*regulatory sites by employing their MYB2 domain but likely could not activate transcription being unable to heteromerize with other transcription factors (because these LFG proteins lacked the protein-binding MYB1 domain). LFG proteins have a putative DNA binding CCHC-type zinc finger domain that likely functions as a generic DNA binder. More specific DNA binding properties come from the MYB2 domain of LFG proteins. The LFG domain and the EAR domain repress transcription of the downstream gene. Phosphorylation of the conserved serine is necessary for interaction with 14-3-3 proteins; in fact, LFG proteins from rice and *Glycine max* have been shown to interact with 14-3-3 proteins for correct localization (Dhaubhadel & Li, 2010; Chen *et al*., 2019).

### DRIF proteins are ancient

DRIF genes were originally described from flowering plants (Machemer *et al*., 2011; Raimundo *et al*., 2013) but have since been reported from across Viridiplantae (Raimundo *et al*., 2018). DRIF proteins—at least in flowering plants—are characterized by a divergent MYB domain that does not always have the same conservation as in other, more typical MYB domains (Raimundo *et al*., 2013). These proteins also have a second conserved region near the C-terminus called the DUF3755. We found homologous, *DRIF-like* genes in the genomes of distantly related eukaryotes—from Glaucophyta, Unikonta, and the SAR clade (Fig. 2). These proteins have regions with homology to the MYBD domain and display limited sequence conservation in the region corresponding to the DUF3755. Some of the DRIF proteins from *Entamoeba histolytica* (Unikonta) are divergent but have still been treated by PANTHER as a homolog of DRIF proteins (‘PTHR14000 in PANTHER’). The DRIF protein in the Glaucophyta species *Cyanophora paradoxa* has a MYBA domain upstream near the N-terminus, in addition to the MYBD domain (Supplementary Fig. 3). We did not find any other gene with a similar *MYBA– MYBD–DUF3755* configuration in any other species, suggesting that the presence of the MYBA domain in a DRIF protein is unique to this branch of the phylogenetic tree. It is important to note that the closest relative of the MYBA domain from *Cyanophora paradoxa* DRIF-like1 is a MYBA domain from a stand-alone MYBA–MYB1 protein from a Oomycota (p=0.88, Supplementary Fig. 3), and not the other MYBA domain in its own genome (present in the DIV protein *Cyanophora paradoxa* DIV-like1).

The region corresponding to DUF3755 is less conserved among DRIF homologs than the MYB domain. Using GenBank conserved domain finder tool on distant relatives of AmDRIF (e.g., from *Entamoeba*) does not identify a DUF3755. This is not surprising, because DUF3755 was initially reported from Viridiplantae (Raimundo *et al*., 2013), and all the currently annotated members are from Viridiplantae (InterPro and GenBank pfam12579). However, based on two lines of evidence, we consider the region corresponding to DUF3755 as homologous between Viridiplantae and other eukaryotes despite low sequence identity. First, despite their divergence the region corresponding to DUF3755 and the region immediately flanking it has some conserved residues across eukaryotes. Second, the degree of divergence among various species for this region corresponds to their species-level relationships. This suggests that the conservation reported from DUF3755 in Viridiplantae evolved gradually from similar, homologous sequences.

The gene tree of DRIFs across eukaryotes largely matches the species tree of Eukaryotes—the first two groups to diverge are genes from Unikonta, followed by SAR+Viridiplantae+Glaucophyta (Fig. 5). This suggests the possibility that DRIF genes (that have both the MYBD domain and the DUF3755) are likely ancestral to all or, at least, a large group of eukaryotes.

**Figure 5.**
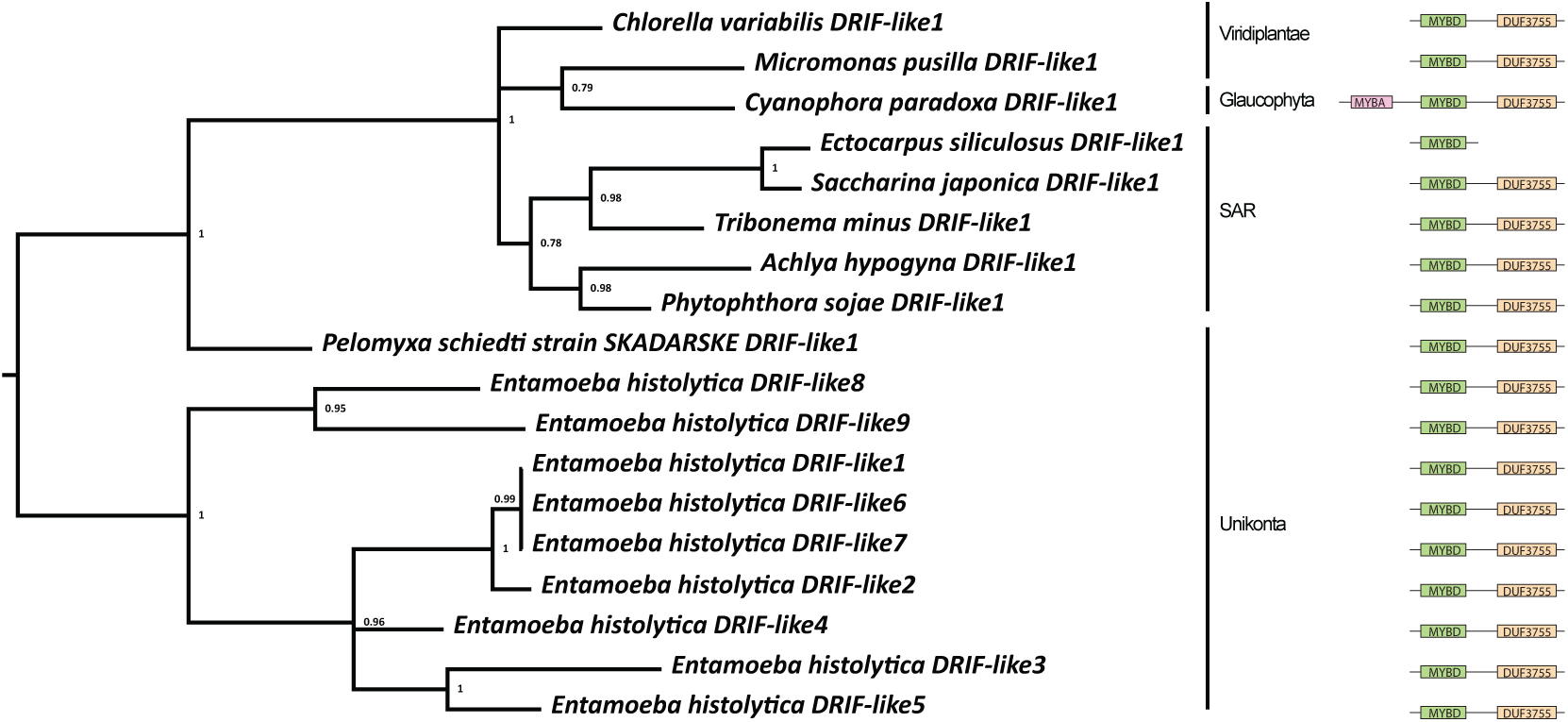
Amino acid-based Bayesian phylogeny of *DRIF*. The tree was rooted at the midpoint. Bayesian posterior probabilities presented at nodes. Domain organization represented on the right.

### Origin of the MYB domains found in DIV and DRIF

*DIV* genes can have up to three *MYB* domains—*MYBA*, *MYB1*, and *MYB2*—and an *HDI* domain. However, given that most of these genes have at least two *MYB* domains, the origin of the individual *MYB* domains has remained unresolved. To understand where these *MYB* domains came from and what other domains are their closest relatives, we reconstructed an amino acid-based Bayesian phylogeny of most MYB domains from representative Phaeophyta species (identified by Zeng *et al*., 2022) along with the MYBA, MYB1, and MYB2 domains from representative eukaryotes (Supplementary Fig. 2). We also reconstructed a smaller tree of the SHAQKY clade to better understand the relationship within the group (Supplementary Fig. 4).

We found that MYBA and MYB2 domains are a part of the larger SHAQKY clade of MYB domains, having the amino acid residues S-H-A-Q-K-Y (or a similar sequence). MYB2 domains in proteins from *DIV* and stand-alone *MYB2–HDI* genes are associated with an HDI domain. These HDI-associated MYB2 domains are nested within SHAQKY domains that are not associated with an HDI domain (Supplementary Fig. 2, Supplementary Fig. 4).

MYBA domains are also members of the SHAQKY clade. However, the corresponding positions in MYBA domains do not have high sequence identity with the residues S-H-A-Q-K-Y. This region is variable even among MYBA domains (Supplementary Fig. 4). It is likely that S-H-A-Q-K-Y or a similar sequence is ancestral to the SHAQKY clade (Supplementary Fig. 4, Supplementary Fig. 2). This would suggest that MYBA domains evolved by substantial divergence from the ancestral gene of the SHAQKY clade.

The origin of MYB1 and the MYBD has been less obvious with no clear sister clades to MYB1 domains (Supplementary Fig. 2), and the divergent MYBD domain (found only in DRIF) was not included in initial analyses. A close relationship between MYB1 domains and the MYBD domains has been suggested but such nodes had poor bootstrap support of 30–39% (Raimundo *et al*., 2018). There are no obvious orthologs of MYB1 domains outside Gyrista and Archaeplastida+Cryptophyta, suggesting that either the branch leading to MYB1 domains or its sister has undergone significant divergence. Therefore, we performed PSI-BLAST (Position-Specific Iterated BLAST) searches in GenBank against selected eukaryotic lineages with a MYB1 domain as the query. PSI-BLAST is designed to identify distant homologs. We reconstructed a phylogeny of MYB domains with these PSI-BLAST hits identified from NCBI, some additional MYB homologs, MYBD domains from manually curated DRIF proteins along with most of the MYB domains used in Supplementary Fig. 2.

We found that MYB1 and the MYBD cluster together with MYB domains from the Zuotin/Zuotin-related factor (ZUO1/ZRF) proteins (Supplementary Fig. 5), particularly the ones classified as the DNAJ HOMOLOG SUBFAMILY C MEMBER 2 (DNAJC2) and the ZINC FINGER, ZZ DOMAIN CONTAINING 3 (ZZZ3). DNAJC2 and ZZZ3 members often have one or two MYB domains (Supplementary Fig. 6)—these domains have been called SANT1 and SANT2, and they are close paralogs of each other (Chen *et al*., 2014)

We reconstructed a phylogeny of MYB1 and MYBD domains with the SANT1/2 domains from DNAJC2 and ZZZ3 proteins (identified in Supplementary Fig. 5). MYB1 domains and the MYBD domains from DRIF proteins are more closely related to SANT2 domains than to SANT1 domains (Bayesian posterior probability = 1, Supplementary Fig. 6), though is not clear whether MYBD and MYB1 are sister to SANT2 or are nested within it (moderate support for monophyly of SANT2 relative to MYB1 or MYBD).

MYB1 domains are found only in Gyrista and Archaeplastida+Cryptophyta, but the following line of evidence suggests that they evolved in the common ancestor or all/most eukaryotes and were lost in several lineages. In Supplementary Fig. 6, the lineage sister to MYB1 domains are the MYB domains from ZZZ3+DRIF proteins—both ZZZ3 and DRIF are found in a wide diversity of eukaryotes suggesting that they evolved near the base of eukaryotes. Further, this clade containing MYB1 domains, and the MYB domains from ZZZ3+DRIF proteins is sister to the SANT2 domains. SANT2 domains are present in all eukaryotes, except fungi, (Chen *et al*., 2014; Shrestha *et al*., 2019), suggesting that SANT2 domains evolved at the base of eukaryotes. If that is the case, the lineage that is sister to SANT2 (which is MYB1+DRIF+ZZZ3) should also have had evolved at the base of eukaryotes. This fact that the two nodes closest to the MYB1 domains can be traced back to the base of eukaryotes suggests that MYB1 domains also evolved near the base of eukaryotes. However, among extant eukaryotes, MYB1 domains are only found in Gyrista and in Archaeplastida+Cryptophyta. This would mean that there have been multiple losses of MYB1 domains across various eukaryotic lineages. We hypothesize a situation where *MYBA–MYB1* genes were present at least in the common ancestor of Gyrista and Archaeplastida+Cryptophyta (if not all eukaryotes) but were lost multiple times.

### Ancestral Function of the conserved MYB and HDI domains

The ancestral functions of MYBA, MYB1, MYB2, and MYBD domains are yet unresolved. The SANT2 domain, which are the closest relatives of MYB1 and MYBD domains, can interact with histones in the Chlorophyta member *Volvox* (Pappas & Miller, 2009), and with multiple kinds of proteins in humans (Kroczynska *et al*., 2004, 2005). This is consistent with the protein-protein interaction displayed by MYB1 and MYBD domain from DRIF in flowering plants.

The ancestral function of the MYB2 domain can be hypothesised based on the common function of proteins with this domain in model plants where it binds to *cis*-regulatory sites of protein coding genes. In many green plants, ancestral stand-alone MYB2 proteins (which evolved before any other MYB2 containing protein with additional MYBA, MYB1 or HDI domains) function in circadian responses (Corellou *et al*., 2009; Nguyen & Lee, 2016). In *Arabidopsis thaliana*, the MYB2 domain is found in typical DIV proteins, LFG proteins, and stand-alone MYB2 proteins. Stand-alone MYB2 proteins and LFG proteins in *A. thaliana* are associated with circadian clock (reviewed in Nguyen & Lee, 2016) and photomorphogenesis (Nguyen *et al*., 2015; Huang *et al*., 2015). We hypothesise that the ancestral function of LFG proteins and DIV proteins (and the corresponding MYB2 domain) was associated with circadian or seasonal photoperiodism. It is then not surprizing that DIV proteins were later co-opted for flower development—a phenotype strongly associated with photoperiodic change.

The N-terminal region of the HDI domain is rich in arginine and lysine. This region is a nuclear localization signal in the rice LFG protein MYBS2 (Chen *et al*., 2019). We hypothesize that the ancestral function of the HDI domain is likely a nuclear localization signal not only in this LFG protein but also in other DIV homologs with HDI domains.

MYB2 and MYBA domains are nested within the SHAQKY clade that is distributed across eukaryotes. In the social slime mold *Dictyostelium discoideum*, a SHAQKY clade protein is responsible for development of the pre-stalk associated with the transition to the reproductive stage (Fukuzawa *et al*., 2006). In *Entamoeba histolytica*, a SHAQKY clade protein is associated with shifts towards a cyst-like stage (as opposed to a trophozoite stage). This putatively ancestral feature of SHAQKY domains in being useful in transcriptionally regulating developmental transitions could possibly have been co-opted in flower development

## Conclusions

We have dissected the origin of the MYB domains found within the DIV-based regulatory networks DIV— including the history of domain gain, domain loss, and gene fusion (Fig. 6). Our data suggest that an ancestral gene with a *MYB2* domain from the *SHAQKY* clade acquired an *HDI* domain, thus becoming a stand-alone *MYB2*–*HDI* gene. This latter gene with the configuration *MYB2–HDI* then fused with a gene with the configuration *MYBA–MYB1*. MYBA domains are also members of the SHAQKY clade. On the other hand, MYB1 and MYBD domains are related to the SANT2 domains of DNAJC2 and ZZZ3 proteins.

**Figure 6.**
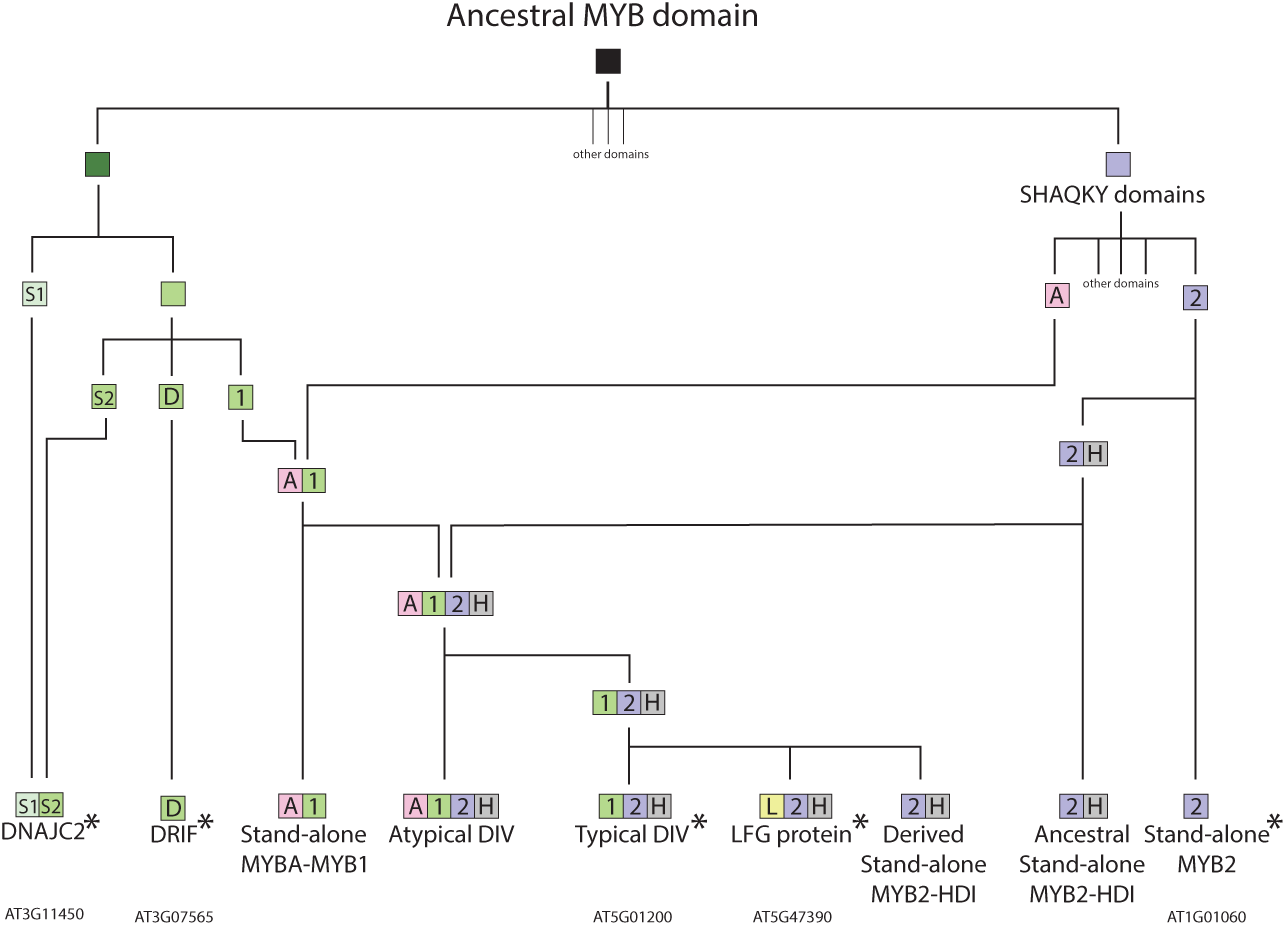
A model representing the origin of the domain configuration associated with DIV and DRIF proteins, and their close homologs. A: MYBA domain; D: MYB domain of DRIF; H: HDI domain; L: LFG domain; S1: SANT1 domain; S2: SANT2 domain; 1: MYB1 domain; 2: MYB2 domain. Proteins with asterisks are present in *Arabidopsis thaliana*, and examples of such proteins from *A. thaliana* are provided under the protein type.

Two distinct DIV-based regulatory networks—in *A. majus*, *S. lycopersicum*, and *O. sativa*—act as on/off switches. One switch involves DIV and RAD, the other switch involves DIV and LFG. No DIV-RAD switch has been tested in *O. sativa* (though it is hypothesized to be ancestral to all angiosperms (Sengupta & Hileman, 2022)). Similarly, no DIV-LFG switch has been tested in *A. majus* or *S. lycopersicum*, though transcriptional repression by LFG proteins is known in *S. lycopersicum* (Liu *et al*., 2024) and *A. thaliana* (Nguyen *et al*., 2015; Huang *et al*., 2015).

We hypothesise that these two switches may constitute a larger three-component switch in seed plants. In such a scenario, the functions of the two MYB domains of the typical DIV proteins may get switched off by two different competitors: the MYB1 domain is switched off by RAD, and the MYB2 domain is switched off by LFG. The DIV-LFG switch involving 14-3-3 proteins is possibly more ancient than flowering plants, and likely evolved near the base of Streptophyta (Supplementary Fig. 7); the DIV-RAD switch would have been added near the base of seed plants (after the evolution of RAD).This recurrent evolution of truncated competitive inhibitors from DIV homologs was likely facilitated by the fact that many DIV proteins have multiple MYB domains, each with a distinct function. Loss of any of the MYB domains in one such multi-domain DIV can generate a competitive inhibitor that inhibits its intact paralogs, thus creating a pair of proteins that act as on/off switches for the two contrasting phenotypes. It is possible that reduction in protein/mRNA sequence length after domain loss facilitates cell-to-cell migration and increases effectives of the competitive interaction. The mRNA of the *LFG* gene *AT5G47390* from *A. thaliana* (Thieme *et al*., 2015) and possibly the protein of *A. majus RADIALIS* can migrate cell-to-cell (Corley *et al*., 2005). In flowering plants, the MYBD domain of DRIF interacts with the MYB1 domain of DIV, and the MYB2 domain of DIV acts as a transcription factor (Machemer *et al*., 2011; Raimundo *et al*., 2013). DRIF proteins are found across a wide diversity of eukaryotes, including Gyrista (Fig. 5), suggesting that the common ancestor of all or most eukaryotes had DRIF proteins, and likely had similar MYB domain interactions (Supplementary Fig. 7). DIV proteins (typical and atypical) are unique to Archaeplastida+Cryptophyta, but proteins with the configuration MYBA–MYB1 are present in Gyrista (Fig. 2). We have provided preliminary evidence that stand-alone MYBA–MYB1 proteins may have been present in the last common ancestor of Gyrista and flowering plants (if not a larger group of eukaryotes). Stand-alone MYBA–MYB1 proteins possibly have similar molecular properties as a typical DIV protein—both have a MYB1 domain, and the other MYB domain (MYBA or MYB2) is from the SHAQKY clade. Given this, we hypothesize a molecular interaction in Gyrista that is similar to the DIV-DRIF interaction in flowering plants: the MYBD domain of DRIF interacting with the MYB1 domain of stand-alone MYBA–MYB1 protein, and the MYBA domain acting as transcription factor (Supplementary Fig. 7). This would suggest that the DIV-DRIF interaction reported in flowering plants are a modification of an ancient genetic network that pre-dates major diversification events of eukaryotes.

## Materials and methods

### Homolog identification

We studied paralogs of four MYB domains in this work—the MYB domains from DRIF proteins (MYBD domain), and the MYB domains from DIV proteins and their homologs (MYBA, MYB1, and MYB2 domains) (Fig. 1 and 2). Additionally, we studied the DUF3755 that is associated with MYBD domains, and the HDI domain that is associated with MYB2 domains.

We identified close homologs of DIV and DRIF by performing protein- and nucleotide-based BLAST searches (Altschul *et al*., 1990) against published databases (Supplementary Tables 1-6). We first used BLAST to search genomes of Viridiplantae and their closest relatives. We used known DIV and DRIF proteins as the query for these BLAST searches—and as we identified more early-diverging homologs, we employed them as queries to search all major clades of Eukaryotes. We searched each major eukaryotic clade in GenBank and by downloading genome sequences of representative taxa from other sources. The distribution of these genes is patchy; therefore, when we detected a homolog in one clade of eukaryotes, we surveyed more species from that clade to generate a denser sampling. We maintained the coding sequence boundaries reported by previous authors, except for *Cryptophyceae sp. CCMP2293 DIV-like1.* The coding sequence of *Cryptophyceae sp. CCMP2293 DIV-like1* was determined based on the EST file that reported a single transcript for this region (Locus8164v1rpkm21.15). However, in the genome file, this locus is split between two closely placed genes (jgi.p|Crypto2293_1|3631476 and jgi.p|Crypto2293_1|2370005). The homologous region in another Cryptophyta species, *Guillardia theta,* has a single gene, consistent with the EST data for *Cryptophyceae sp. CCMP2293 DIV-like1*. Therefore, we treated *Cryptophyceae sp. CCMP2293 DIV-like1* as a single gene as reported in the EST file.

We identified most MYB domains in representative Phaeophyta (Brown Algae) species by searching for conserved MYB domains in a list of MYB proteins identified previously (Zeng *et al*., 2022). The list by Zeng *et al.,* (2022), is not exhaustive but is sufficient to generate a backbone phylogeny of MYB domains. To predict the MYB domains in these proteins (Zeng *et al*., 2022), we implemented Interproscan (Blum *et al*., 2020) in Geneious (Kearse *et al*., 2012). It is possible that highly divergent MYB domains may have remained unidentified in our search or in the search implemented by Zeng *et al*. (Zeng *et al*., 2022). However, we used previously known MYBA, MYB1, and MYB2 domains, as queries to search the list of Phaeophyta MYB proteins, thus ensuring that the close relatives of each domain were identified.

### Phylogenetic analysis

We estimated the relationships among various MYB genes or MYB domains by employing MrBayes (Ronquist *et al*., 2012) available at CIPRES (Miller *et al*., 2010). We estimated the phylogenetic relationships among genes with *MYB2* and *HDI* domains (Fig. 3) based on a nucleotide alignment. We performed an alignment with the MAFFT alignment tool (Katoh *et al*., 2002) implemented in Geneious (Kearse *et al*., 2012) and manually refined the alignment. We then employed minimally informative priors for MrBayes analysis and rooted the resulting 50%-majority rule consensus tree at the midpoint (Fig. 3). We estimated the phylogenetic relationships among genes with *MYBA* and/or *MYB1* domains (Fig. 4) with an approach similar to those with *MYB2–HDI* domains.

We determined the relationship of MYBA, MYB1, and MYB2 domains with other MYB domains by reconstructing a tree of most MYB domains from representative Phaeophyta species along with MYBA, MYB1, and MYB2 domains from representative species. We focused on Phaeophyta species because these are among the most distant lineages (relative to Viridiplantae) that have MYBA and MYB1 domains in their genomes (MYB2 domains are present more widely). This tree of most MYB domains (Supplementary Fig. 2) has the following taxa: all MYB domains identified from representative Phaeophyta species (*Ectocarpus siliculosus*, *Nemacystus decipiens*, *Cladosiphon okamuranus*, *Saccharina japonica*); MYBA domains from Cryptophyta and Glaucophyta DIV proteins; MYB1 domains from Cryptophyta, Glaucophyta, and Chlorophyta; and MYB2 domains associated with HDI domains from representative species across eukaryotes. We estimated the phylogenetic relationships among these MYB domains based on an amino acid alignment. We performed an alignment with the MAFFT alignment tool (Katoh *et al*., 2002) implemented in Geneious (Kearse *et al*., 2012) and manually refined the alignment. We then employed minimally informative priors for MrBayes analysis and rooted the resulting 50%-majority rule consensus tree at the midpoint (Supplementary Fig. 2).

To better understand the relationship among the SHAKQY clade of MYB domains (that includes MYBA and MYB2 domains), we reconstructed a tree of the SHAKQY clade (Supplementary Fig. 4) using a subset of genes from Supplementary Fig. 2. We reduced the number of Phaeophyta from four species to two in this alignment to minimize polytomies. Alignment and Bayesian phylogeny was performed on a translated alignment as in Supplementary Fig. 2.

We identified *DRIF* genes from across eukaryotes by BLAST searches. We estimated the phylogenetic relationships among *DRIF* genes (Fig. 5) based on a translated alignment. We performed an alignment with the MAFFT alignment tool (Katoh *et al*., 2002) implemented in Geneious (Kearse *et al*., 2012) and manually refined the alignment. We then employed minimally informative priors for MrBayes analysis and rooted the resulting 50%-majority rule consensus tree at the midpoint (Fig. 5).

The relationship between MYB1 domains with other MYB domains, especially, the MYBD domain from DRIF proteins, was not clear from Supplementary Fig. 2 because the list of MYB proteins from Zeng et al. (Zeng *et al*., 2022) did not include DRIFs. A closer relationship between MYB1 domains and MYBD domains has been suggested by other workers (Raimundo *et al*., 2018). Therefore, we reconstructed their phylogenetic history using the following sets of MYB domains. *Set-1*: BLAST identified potential relatives of MYB1 from diverse eukaryotes. We used the MYB1 domain from *Saprolegnia parasitica* MYBgene1 as the query and performed two separate PSI-BLAST (Position-Specific Iterated BLAST) searches in GenBank against the Dictyostelid Cellular Slime Molds (which are a clade within of Unikonta, taxid:33083) and SAR supergroup (taxid:2698737; excluding Oomycota and Ochrophyta because we already had MYB domains from them). We confirmed that the resulting hits were MYB domains by testing them with Interproscan. MYB1 domains were only found in one subclade of the SAR clade and in Archaeplastida+Cryptophyta suggesting the possibility that the MYB1 domains in the close relatives may have diverged beyond identification by simple alignment or regular BLAST. We used PSI-BLAST to better capture such divergent domains. We know that this approach of identifying divergent MYB1 domains was successful because in the subsequent phylogeny (Supplementary Fig. 5) both close and distant homologs of MYB1 domains were identified. This means that PSI-BLAST had exhausted identifying close (but divergent) homologs of the MYB1 domain before moving on to identifying distant homologs. *Set-2*: additional homologs of some of the hits identified in set-1. These domains are from the POWERDRESS clade of proteins. *Set-3*: MYBD domains from DRIF homologs identified from across eukaryotes by BLAST searches. *Set-4*: the MYB domains used in Supplementary Fig. 2, but the sequences from some of the Phaeophyta species were removed (to create a more manageable computational dataset). We estimated the phylogenetic relationships among these MYB domains from sets 1–4 based on an amino acid alignment. We performed an alignment with the MAFFT alignment tool (Katoh *et al*., 2002) implemented in Geneious (Kearse *et al*., 2012) and manually refined the alignment. We then employed minimally informative priors for MrBayes analysis and rooted the resulting 50%-majority rule consensus tree at the midpoint (Supplementary Fig. 5).

From Supplementary Fig. 5, we identified the clade that included MYB1 domains and the MYBD domains from DRIF proteins. This clade has MYB domains from Zuotin/Zuotin-related factor (ZUO1/ZRF) proteins (Supplementary Fig. 5), particularly the ones classified as the DNAJ HOMOLOG SUBFAMILY C MEMBER 2 (DNAJC2) and the ZINC FINGER, ZZ DOMAIN CONTAINING 3 (ZZZ3). These proteins have two kinds of MYB domains—SANT1 and SANT2, sometimes in the same protein. Our PSI-BLAST search had identified a non-exhaustive set of SANT1/2 domains for the ZUO1/ZRF homologs captured by it. To better understand the relationship of MYB1 and MYBD with SANT1/2 domains, we downloaded all SANT1/2 homologs from these proteins. We then reconstructed a phylogeny of these domains along with MYB1 and MYBD domains. We generated an alignment and reconstructed a phylogenetic tree as described before. We rooted the resulting 50%-majority rule consensus tree at the midpoint (Supplementary Fig. 6). The sequences of the genes and proteins used in these analyses, their sources, and the alignment files (including the Bayesian command block) associated with the phylogenetic studies are available in the supplementary files.

### Tests of selection

We performed tests of selection using the programme RELAX (Wertheim *et al*., 2015) available online at Datamonkey (‘RELAX at Datamonkey’). RELAX can detect whether a subset of branches in a phylogenetic tree (test branches) is undergoing relaxed vs. intensification of selection relative to another set of branches (reference branches). We tested whether the *MYBA* domains from post-fusion *DIV* genes (and their descendants) are undergoing relaxed vs. intensification of selection relative to the *MYBA* domains from pre-fusion genes (i.e., *MYBA* domains from genes with only *MYBA*-*MYB1* configuration). In the test of selection on *MYBA* domains, *Micromonas pusilla DIV-like1* was not included in the test group nor the reference group because it has no *MYBA* domain. The test and reference groups are marked in Fig. 4, which we used as the backbone tree. The input file (including the alignment used) is available in Supplementary file 6. For the selection tests on *MYBA* domains, we only used the *MYBA* domain and no flanking sequences (because the flanking regions are poorly conserved).

We performed similar tests of relaxed vs. intensified selection on *MYB1* and *MYB2* domains before and after the gene fusion event. For *MYB1*, the test and reference groups are marked in Fig. 4, which we used as the backbone tree; the input file is available in Supplementary file 7; the input alignment included the *MYB1* domain and some conserved region immediately downstream. For *MYB2,* test and reference groups are marked in Fig. 3, which we used as the backbone tree; the input file is available in Supplementary file 3; the input alignment included the *MYB2* domain and some conserved region immediately upstream and downstream. We performed each test multiple times to ensure that all or the majority of the replicates agreed on the results.

## Acknowledgements

The authors acknowledge the advice from Dr. Joel Wertheim regarding RELAX. The authors thank members of the Howarth lab.

## Author contributions

ASG conceptualized the project and developed it with advice from DH. ASG performed all analyses and prepared the first draft of the manuscript. DH and ASG revised the manuscript. All authors read and approved the manuscript.

## Data availability

The data underlying this article are available in the article and in its online supplementary material.

## Supplemental data

### Supplementary figures

**Supplementary Fig. 1.** A simplified tree of eukaryotes representing the relationship among organisms mentioned in this study. Different parts of this tree were drawn based on published trees from the following sources: backbone (Burki *et al*., 2012), Streptophyta (Wang *et al*., 2020), Stramenopila (Thakur *et al*., 2019), and Land plants (Liu *et al*., 2022).

**Supplementary Fig. 2.** Amino acid-based Bayesian phylogeny of MYB domains from representative Phaeophyta species along with MYBA, MYB1, and MYB2 domains, from representative eukaryotes. Names of MYBA, MYB1, and MYB2 domains start with “MYB_A/1/2_domain” (e.g., **MYB_2_domain**_of_Chara_braunii_DIV_like1_BFEA01000398_SHAQKY). Names of other MYB domains (that are not MYBA, MYB1, or MYB2 domains) start with the species name (e.g., **Ectocarpus_siliculosus**_Ec_06_008620_MYB_domain_2). Note that such domains (that are not MYBA, MYB1, or MYB2 domains) may come from genes with multiple MYB domains and hence these domains are numbered near the end of the name with the suffix “MYB_domain_1/2/3,” etc. (e.g., Ectocarpus_siliculosus_Ec_06_008620_**MYB_domain_2**). For SHAQKY clade MYB domains, the amino acid sequence corresponding to the amino acids SHAQKY (SHAQKY, HHARYH, etc.) is provided at the end of the name (e.g., MYB_2_domain_of_Chara_braunii_DIV_like1_BFEA01000398_**SHAQKY**). The tree was rooted at the midpoint. Bayesian posterior probabilities presented at nodes.

**Supplementary Fig. 3.** Amino acid-based Bayesian phylogeny MYB domains including the two MYB domains from *Cyanophora paradoxa* DRIF-like1. This protein has the expected MYBD domain found in all DRIF proteins (indicated with a red arrow) but also has a MYBA domain (indicated with a blue arrow). Note that *C. paradoxa* also has another MYBA domain in the atypical DIV protein named *C. paradoxa* DIV-like1 (indicated by a green arrow). The tree is based on the sequences used in Supplementary Fig. 5 and the two domains from *C. paradoxa* DRIF-like1. The tree was rooted at the midpoint. Bayesian posterior probabilities presented at nodes.

**Supplementary Fig. 4.** Amino acid-based Bayesian phylogeny of all SHAQKY clade proteins from Phaeophyta species along with MYBA, MYB1, and MYB2 domains, from representative eukaryotes. The taxa are a subsample of Supplementary Fig. 2. The tree was rooted at the midpoint. Bayesian posterior probabilities presented at nodes.

**Supplementary Fig. 5.** Amino acid-based Bayesian phylogeny to identify the closest relatives of MYB1 domains and the MYBD domains from DRIF proteins. The tree includes MYB domains from Supplementary Fig. 2 (some species removed to create a more manageable computational dataset), BLAST hits when Unikonta and SAR lineages in NCBI were searched with MYB1 domain as a query, MYB domains from DRIF proteins, and some manually curated MYB domains. The tree was rooted at the midpoint. Bayesian posterior probabilities presented at nodes.

**Supplementary Fig. 6.** Amino acid-based Bayesian phylogeny of MYB1 domains, MYBD domains from DRIFs, and their closest relatives. Each terminal taxon is a MYB domain. The configuration of the corresponding protein is represented as a cartoon. Note that some genes have multiple MYB domains represented by different terminal taxa. The MYB domain at a given tip is represented by a red-bordered box in the cartoon of the protein. The tree was rooted at the midpoint. Bayesian posterior probabilities presented at nodes.

**Supplementary Fig. 7.** Hypothetical step-wise evolution of the molecular interactions of DIV homologs in eukaryotes. Known or hypothesized molecular interactions in representative extant species on the right, putative ancestral states of these interactions at major nodes on the left. Relationships among the clades to which these species belong are represented by a simplified phylogeny of eukaryotes (adapted from Burki *et al*., 2012), with an added branch for extinct hypothetical eukaryotes. Domain types are similar to those in Fig. 2; with additional representation of 14-3-3 proteins, proteins with SANT1/2 domains, and hypothesized DNA binding domains. Dashed lines represent hypothesized molecular interactions. Molecular interactions were hypothesized based on experimental evidence from homologous proteins/domains as follows: SHAQKY clade domains (MYBA, MYB2, and MYB2* domains) binding to DNA (Rose *et al*., 1999; Raimundo *et al*., 2013), SANT1/2 domains interacting with diverse proteins (Kroczynska *et al*., 2004, 2005; Pappas & Miller, 2009; Weaver *et al*., 2019), MYBD domains binding to MYB1 domains (Machemer *et al*., 2011; Raimundo *et al*., 2013; Petzold *et al*., 2018), truncated DIV homologs lacking the MYB2 (i.e., RADIALIS) competing with intact typical DIV proteins for MYD domains (Machemer *et al*., 2011; Raimundo *et al*., 2013), truncated DIV homologs lacking the MYB1 (LFG proteins or their close relatives in Chlorophyta) competing with intact typical DIV proteins for DNA (Chen *et al*., 2019), and LFG proteins interacting with 14-3-3 proteins (Dhaubhadel & Li, 2010; Li & Dhaubhadel, 2012; Chen *et al*., 2019)

### Supplementary files

**Supplementary file 1.** Nexus alignment and MrBayes command block used in Fig. 3 (tree of genes with MYB2–HDI domains).

**Supplementary file 2.** Unaligned coding sequences of genes used in Fig. 3 (tree of genes with MYB2-HDI domains).

**Supplementary file 3.** Input file for RELAX test associated with Fig. 3 (tree of genes with MYB2-HDI domains).

**Supplementary file 4.** Nexus alignment and MrBayes command block used in Fig. 4 (tree of genes with MYBA–MYB1 domains).

**Supplementary file 5.** Unaligned coding sequences of genes used in Fig. 4 (tree of genes with MYBA–MYB1 domains).

**Supplementary file 6.** Input file for RELAX test associated with MYBA domains of Fig. 4 (tree of genes with MYBA–MYB1 domains).

**Supplementary file 7.** Input file for RELAX test associated with MYB1 domains of Fig. 4 (tree of genes with MYBA–MYB1 domains).

**Supplementary file 8**. Nexus alignment and MrBayes command block used in Supplementary Fig. 2 (tree of MYB domains).

**Supplementary file 9.** Unaligned sequences used in Supplementary Fig. 2 (tree of MYB domains).

**Supplementary file 10**. Nexus alignment and MrBayes command block used in Supplementary Fig. 4 (tree of SHAQKY domains).

**Supplementary file 11.** Unaligned coding sequences of *DRIF* genes used in Fig. 5.

**Supplementary file 12.** Nexus alignment and MrBayes command block used in Fig. 5 (tree of *DRIF* genes).

**Supplementary file 13.** Nexus alignment and MrBayes command block used in Supplementary Fig. 5 (tree of MYB domains with focus on MYB1 and DRIF).

**Supplementary file 14.** Nexus alignment and MrBayes command block used in Supplementary Fig. 6 (tree of MYB1, DRIF, and close relatives).

**Supplementary file 15.** Nexus alignment and MrBayes command block used in Supplementary Fig. 3 (MYB domains with MYBA from *Cyanophora paradoxa* DRIF-like1)

### Supplementary tables

**Supplementary table 1.** Sources of genes used in Fig. 3 (tree of genes with MYB2–HDI domains)

**Supplementary table 2.** Sources of genes used in Fig. 4 (tree of genes with MYBA–MYB1 domains).

**Supplementary table 3.** Source of genes used in Supplementary Fig. 2 (tree of MYB domains).

**Supplementary table 4.** Source of genes used in Fig. 5 (tree of DRIF genes).

**Supplementary table 5.** Source of MYB domains identified by performing PSI-BLAST with MYB1 domain as a query. All entries are from GenBank.

**Supplementary table 6.** Source of POWERDRESS proteins.

