## Supplementary material for "On/off switches in the *DIVARICATA*-based regulatory network evolved through gene duplication, fusion, and truncation": Supplementary figures.pdf

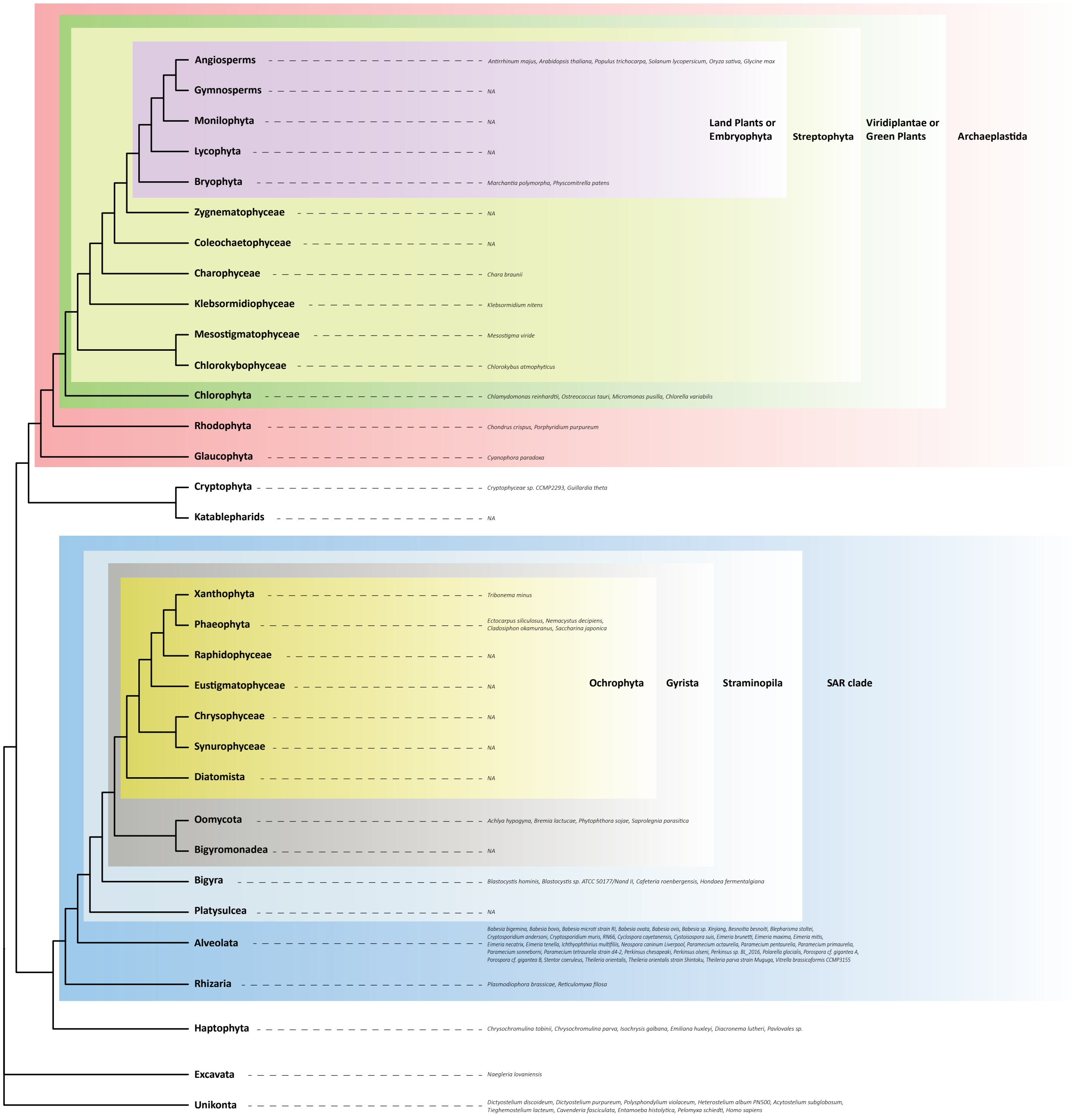


Supplementary Fig. 1. A simplified tree of eukaryotes representing the relationship among organisms mentioned in this study. Different parts of this tree were drawn based on published trees from the following sources: backbone (Burki et al., 2012), Streptophyta (Wang et al., 2020), Stramenopila (Thakur et al., 2019), and Land plants (Liu et al., 2022)

Supplementary Fig. 2. Amino acid-based Bayesian phylogeny of MYB domains from representative Phaeophyta species along with MYBA, MYB1, and MYB2 domains, from representative eukaryotes. Names of MYBA, MYB1, and MYB2 domains start with “MYB_A/1/2_domain” (e.g., **MYB_2**_domain_of_Chara_braunii_DIV_like1_BFEA01000398_SHAQKY). Names of other MYB domains (that are not MYBA, MYB1, or MYB2 domains) start with the species name (e.g., **Ectocarpus_siliculosus**_Ec_06_008620_MYB_domain_2). Note that such domains (that are not MYBA, MYB1, or MYB2 domains) may come from genes with multiple MYB domains and hence these domains are numbered near the end of the name with the suffix “MYB_domain_1/2/3,” etc. (e.g., Ectocarpus_siliculosus_Ec_06_008620_**MYB_domain_2**). For SHAQKY clade MYB domains, the amino acid sequence corresponding to the amino acids SHAQKY (SHAQKY, HHARYH, etc.) is provided at the end of the name (e.g., MYB_2_domain_of_Chara_braunii_DIV_like1_BFEA01000398_**SHAQKY**). The tree was rooted at the midpoint. Bayesian posterior probabilities presented at nodes.

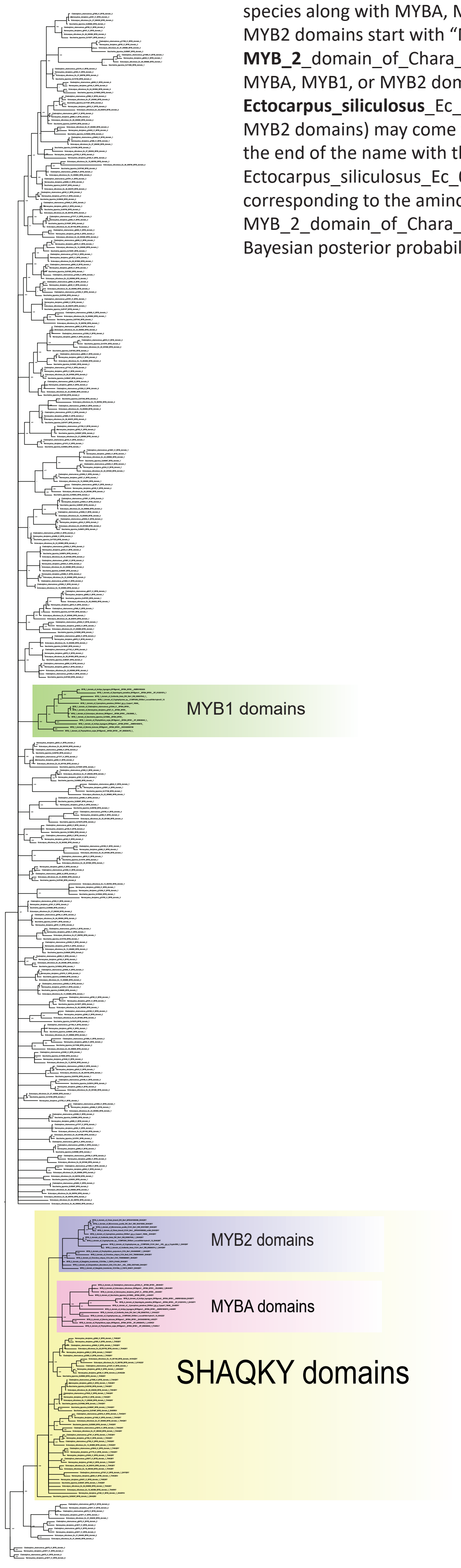

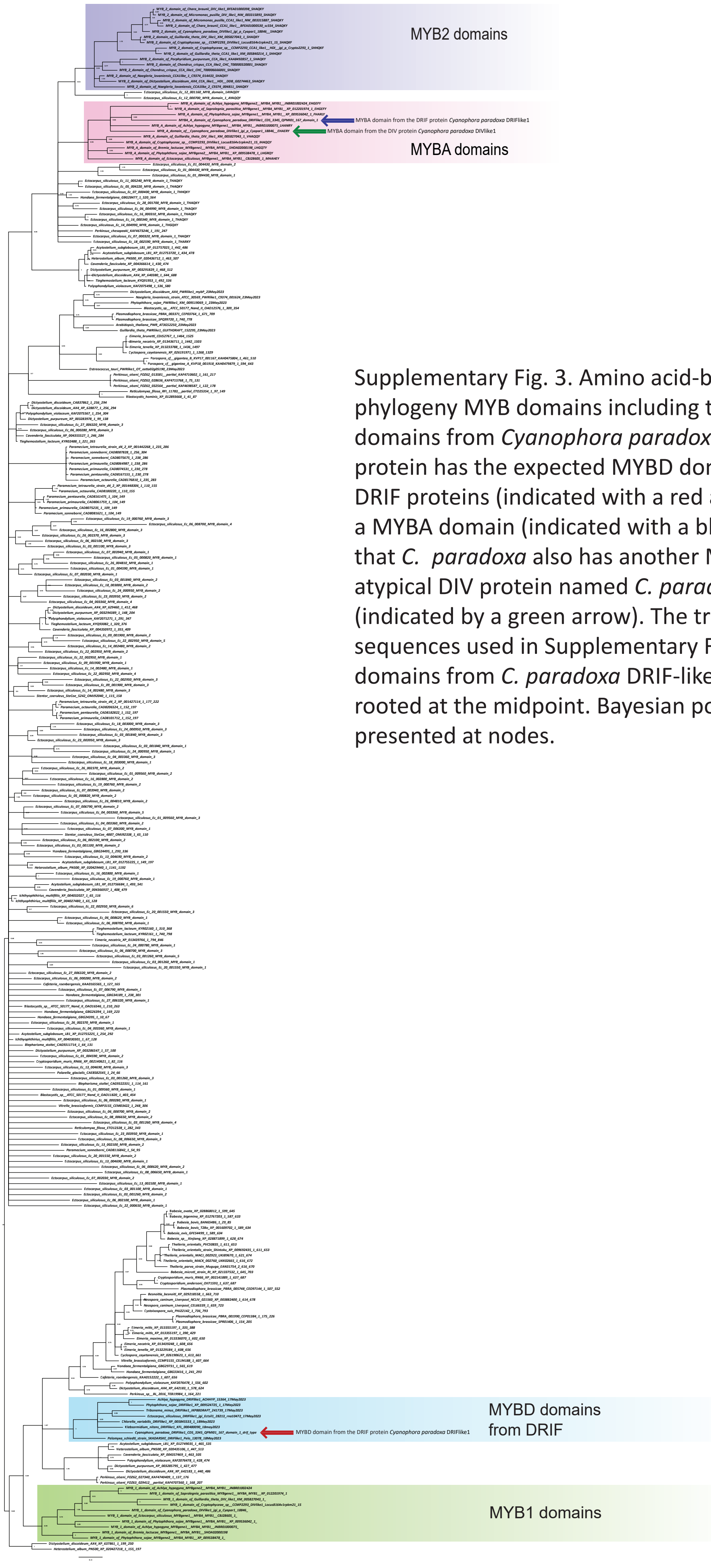


Supplementary Fig. 3. Amino acid-based Bayesian phylogeny MYB domains including the two MYB domains from *Cyanophora paradoxa* DRIF-like1. This protein has the expected MYBD domain found in all DRIF proteins (indicated with a red arrow) but also has a MYBA domain (indicated with a blue arrow). Note that *C. paradoxa* also has another MYBA domain in the atypical DIV protein named *C. paradoxa* DIV-like1 (indicated by a green arrow). The tree is based on the sequences used in Supplementary Fig. 5 and the two domains from *C. paradoxa* DRIF-like1. The tree was rooted at the midpoint. Bayesian posterior probabilities presented at nodes.

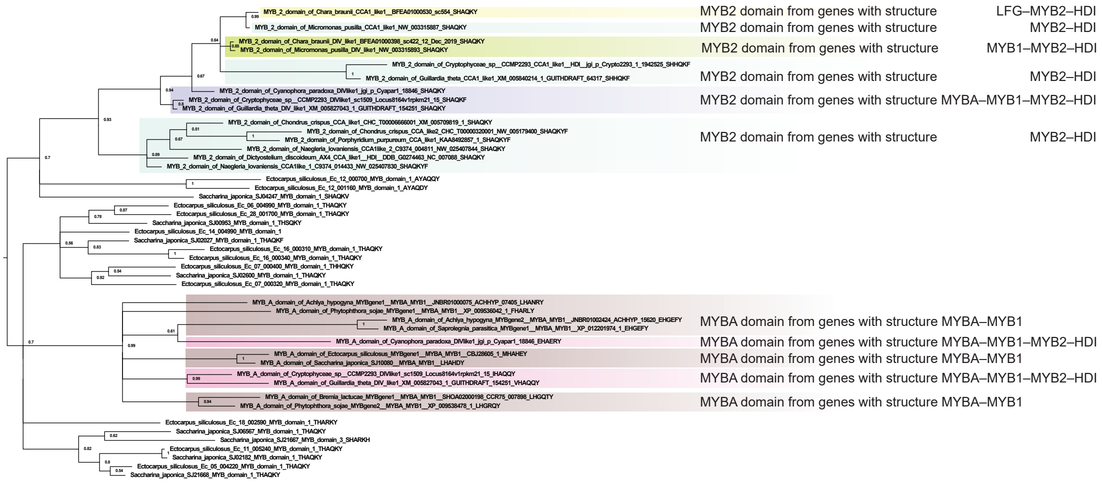


Supplementary Fig. 4. Amino acid-based Bayesian phylogeny of all SHAQKY clade proteins from Phaeophyta species along with MYBA, MYB1, and MYB2 domains, from representative eukaryotes. The taxa are a subsample of Supplementary Fig. 2. The tree was rooted at the mid-point. Bayesian posterior probabilities presented at nodes.

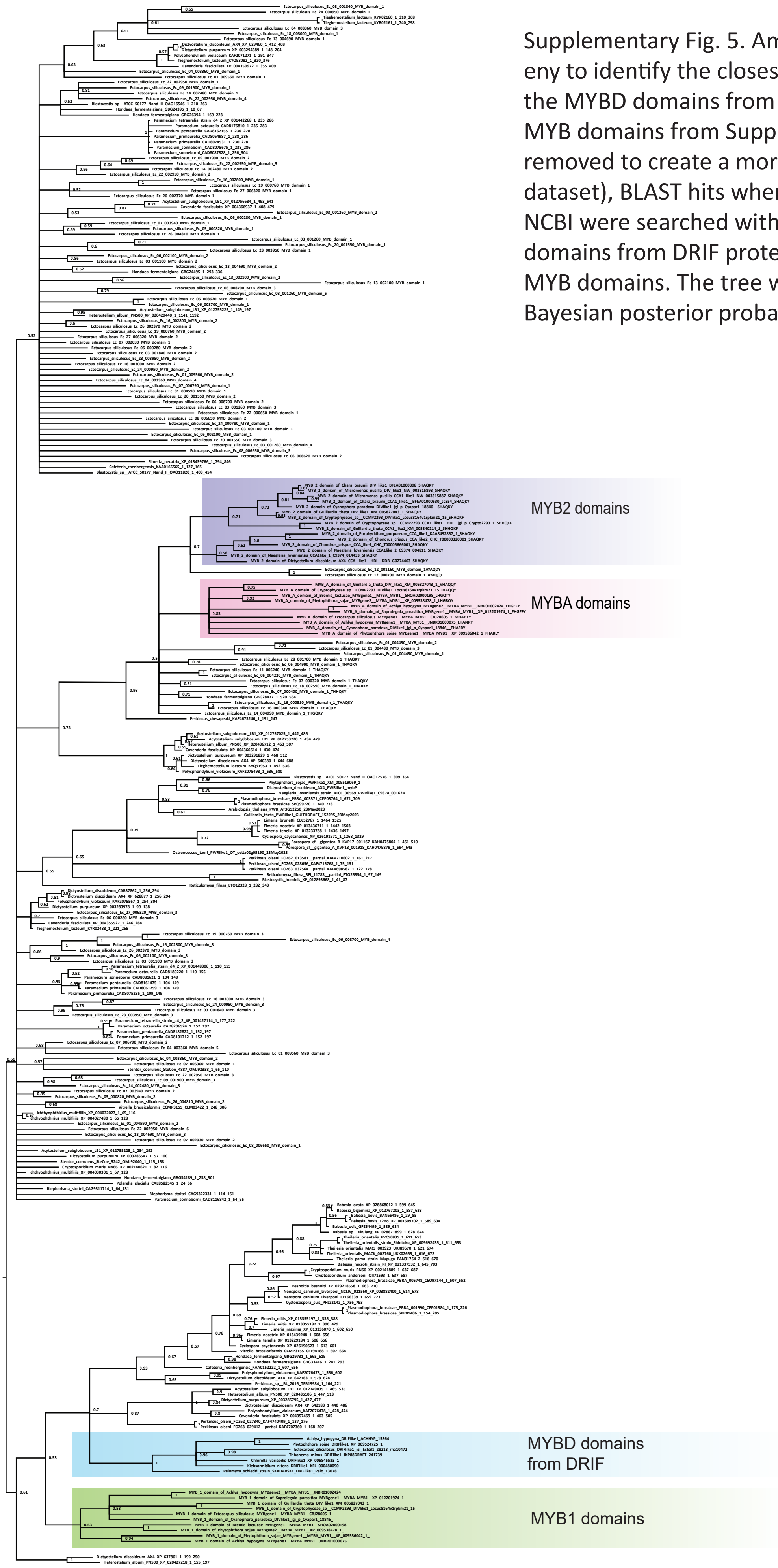


Supplementary Fig. 5. Amino acid-based Bayesian phylogeny to identify the closest relatives of MYB1 domains and the MYBD domains from DRIF proteins. The tree includes MYB domains from Supplementary Fig. 2 (some species removed to create a more manageable computational dataset), BLAST hits when Unikonta and SAR lineages in NCBI were searched with MYB1 domain as a query, MYB domains from DRIF proteins, and some manually curated MYB domains. The tree was rooted at the midpoint. Bayesian posterior probabilities presented at nodes.

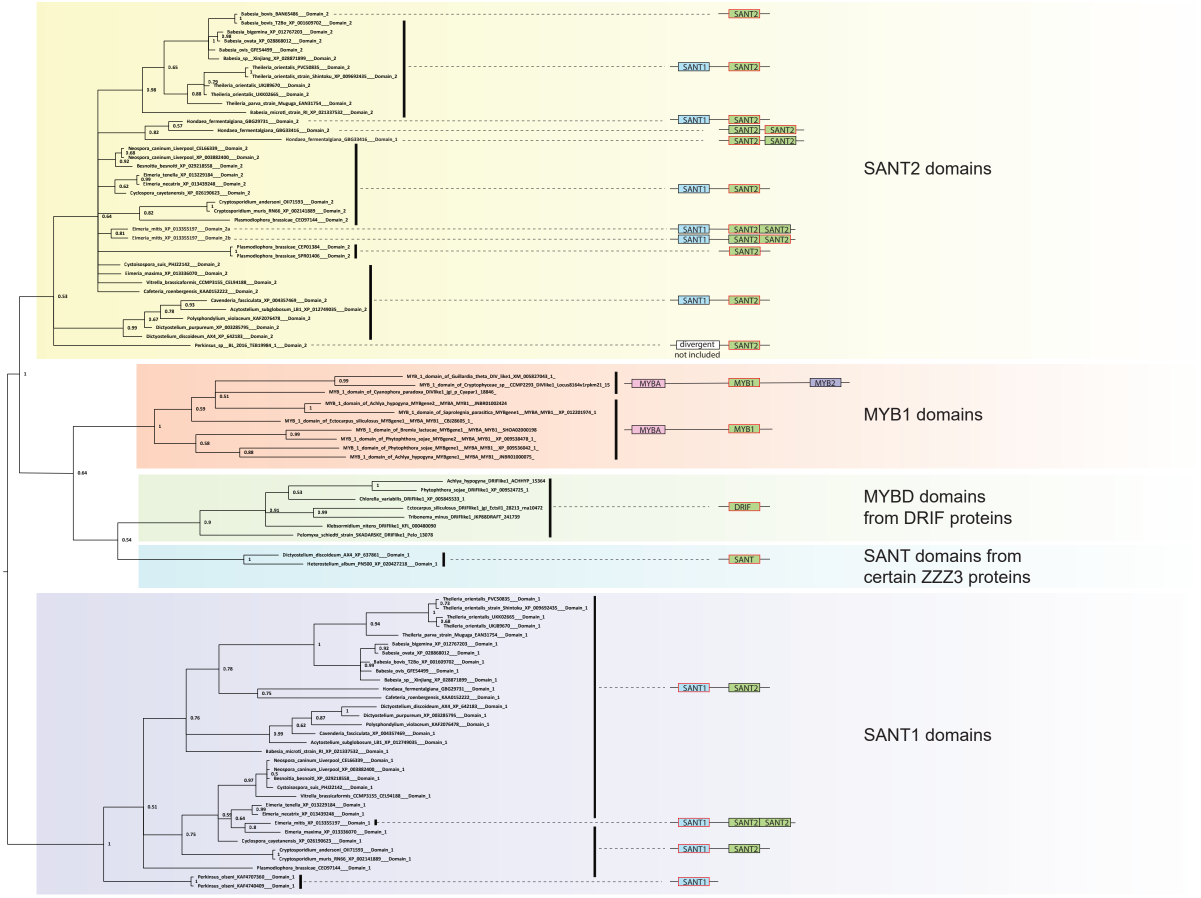


Supplementary Fig. 6. Amino acid-based Bayesian phylogeny of MYB1 domains, MYBD domains from DRIFs, and their closest relatives. Each terminal taxon is a MYB domain. The configuration of the corresponding protein is represented as a cartoon. Note that some genes have multiple MYB domains represented by different terminal taxa. The MYB domain at a given tip is represented by a red-bordered box in the cartoon of the protein. The tree was rooted at the midpoint. Bayesian posterior probabilities presented at nodes.

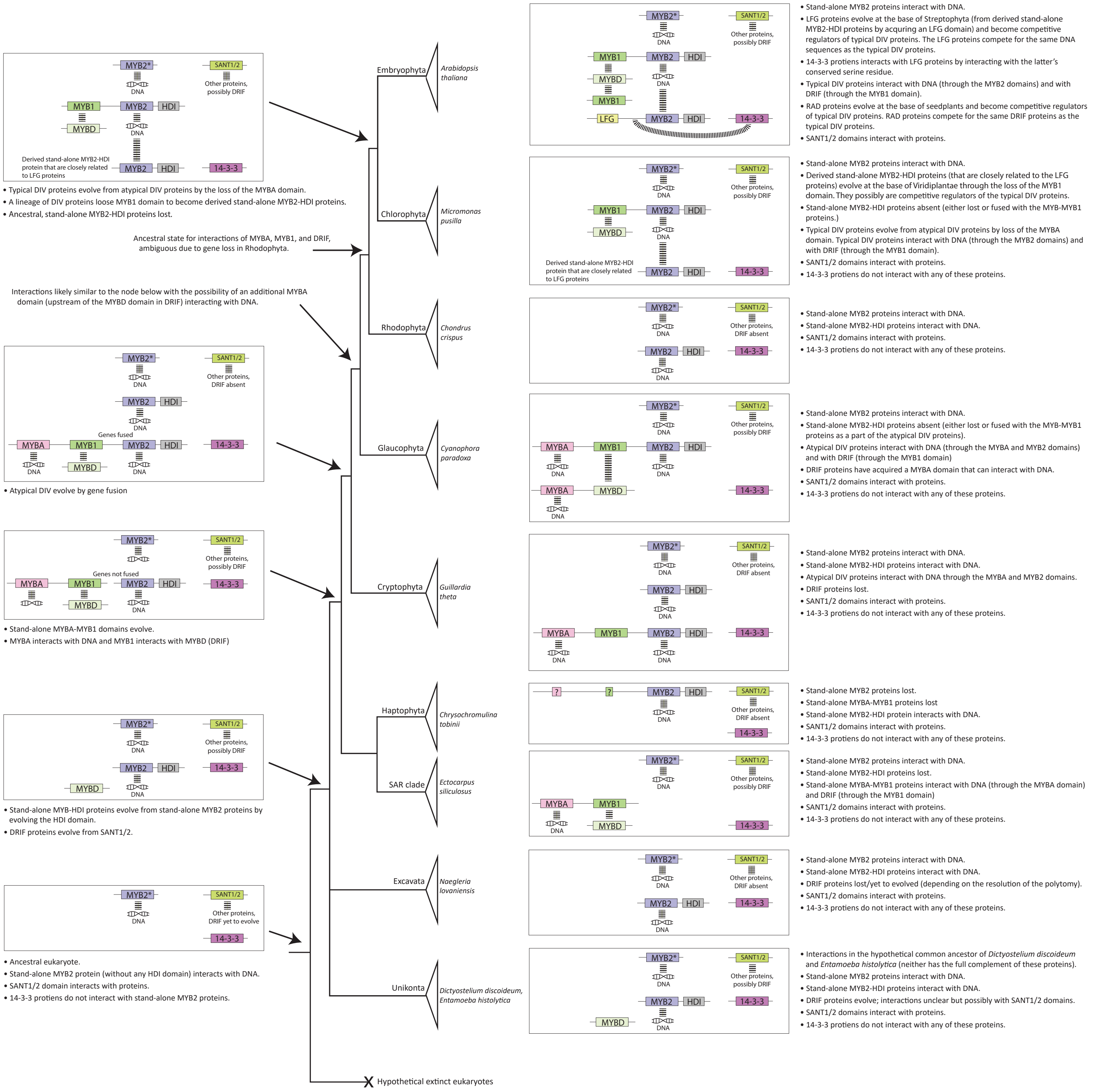


Supplementary Fig. 7. Hypothetical step-wise evolution of the molecular interactions of DIV homologs in eukaryotes. Known or hypothesized molecular interactions in representative extant species on the right, putative ancestral states of these interactions at major nodes on the left. Relationships among the clades to which these species belong are represented by a simplified phylogeny of eukaryotes (adapted from Burki et al., 2012), with an added branch for extinct hypothetical eukaryotes. Domain types are similar to those in Fig. 2; with additional representation of 14-3-3 proteins, proteins with SANT1/2 domains, and hypothesized DNA binding domains. Dashed lines represent hypothesized molecular interactions. Molecular interactions were hypothesized based on experimental evidence from homologous proteins/domains as follows: SHAQKY clade domains (MYBA, MYB2, and MYB2* domains) binding to DNA (Rose et al., 1999; Raimundo et al., 2013), SANT1/2 domains interacting with diverse proteins (Kroczynska et al., 2004, 2005; Pappas & Miller, 2009; Weaver et al., 2019), MYBD domains binding to MYB1 domains (Machemer et al., 2011; Raimundo et al., 2013; Petzold et al., 2018), truncated DIV homologs lacking the MYB2 (i.e., RADIALIS) competing with intact typical DIV proteins for MYD domains (Machemer et al., 2011; Raimundo et al., 2013), truncated DIV homologs lacking the MYB1 (LFG proteins or their close relatives in Chlorophyta) competing with intact typical DIV proteins for DNA (Chen et al., 2019), and LFG proteins interacting with 14-3-3 proteins (Dhaubhadel & Li, 2010; Li & Dhaubhadel, 2012; Chen et al., 2019).
